# A reference map of the human proinsulin biosynthetic interaction network

**DOI:** 10.1101/699132

**Authors:** Duc T. Tran, Anita Pottekat, Saiful A. Mir, Insook Jang, Salvatore Loguercio, Alexandre Rosa Campos, Reyhaneh Lahmy, Ming Liu, Peter Arvan, William E. Balch, Randal J. Kaufman, Pamela Itkin-Ansari

## Abstract

The beta-cell protein synthetic machinery is dedicated to the production of insulin, which plays a critical role in organismal homeostasis. Insulin synthesis requires the proper folding and trafficking of its precursor, proinsulin, yet the precise network of proinsulin protein interactions in the secretory pathway remains poorly defined. In the present study we conducted unbiased profiling of the proinsulin interactome in human islets, utilizing a human proinsulin-specific monoclonal antibody for affinity purification and mass spectrometry. Stringent analysis identified a central node of interactions between human proinsulin and sequential secretory pathway proteins that is remarkably conserved across 3 ethnicities and both genders. Among the most prominent proinsulin interactions was with ER-localized peroxiredoxin-4 (PRDX4). A functional role for PRDX4 in beta-cells was demonstrated by gene silencing that rendered proinsulin susceptible to misfolding, particularly in response to oxidative stress. Conversely, exogenous PRDX4 improved proinsulin folding. Notably, oxidative stress and even high glucose treatment alone induced proinsulin misfolding in human islets and MIN6 cells, and this was accompanied by sulfonylation of PRDX4, a modification known to inactivate peroxiredoxins. This finding prompted PRDX4 analysis in a panel of human islet samples that revealed significantly higher levels of sulfonylated (inactive) PRDX4 in islets from patients with T2D compared to that of healthy individuals. Taken together, these data highlight the importance of elucidating the complete proinsulin interactome in human islets in order to understand critical steps controlling insulin biosynthesis, beta cell function, and T2D.

## Introduction

Globally, an estimated 422 million adults are living with diabetes (http://www.who.int/en/). Moreover, diabetes and its associated complications is a growing cause of death in both developed and developing countries. Beta cell physiology is tightly linked to all forms of diabetes; a better understanding of beta cell physiology is thus necessary to combat this disease.

Insulin biosynthesis begins with translocation of preproinsulin into the endoplasmic reticulum (ER) (1). Following cleavage of the signal peptide, proper proinsulin folding results in a three-dimensional structure stabilized by 3 intramolecular disulfide bonds (1). Proinsulin is then transported into the Golgi complex, encounters zinc, further assembles into hexamers, and is finally transported to immature secretory granules, in which proteolytic processing enzymes cleave proinsulin to mature insulin (and C-peptide) for release in response to glucose (1). Only properly folded proinsulin molecules can transit to the Golgi apparatus from which secretory granules are formed.

Every second, 6000 preproinsulin molecules are delivered to ER for assisted folding (2). Thus, the majority of the beta-cell’s protein synthetic machinery is dedicated to proinsulin production, yet many of the proteins participating in its folding and trafficking are unknown. The critical role of proinsulin’s proper folding is exemplified by its rare heterozygous mutations that prevent correct disulfide bond formation. These misfolded proinsulin molecules from a single mutant allele can entrap wild-type proinsulin in the ER, causing Mutant *INS*-gene-induced Diabetes of Youth (MIDY) both in humans and animal models (1). It is increasingly appreciated that a portion of proinsulin misfolds even in healthy islets (3). We recently showed that this portion of misfolded proinsulin is exacerbated by oxidative stress (4), ER stress (3), or cleavage-inactivation of the chaperone BiP by a specific bacterial protease SubAB (3).

Type I and Type II diabetes are typically associated with inflammation or oxidative stress, which may contribute to proinsulin misfolding (3) thereby limiting the size of the properly-folded proinsulin population. Misfolded proinsulin activates the unfolded protein response (UPR) signaled through two protein kinases PERK and IRE1 and the transcription factor ATF6 (5). Early in progression to diabetes, the three branches of the UPR adapt the beta cell’s ER to accommodate ever-increasing proinsulin synthesis to prevent further accumulation of misfolded proinsulin by (i) attenuating proinsulin synthesis, (ii) inducing expression of genes that promote translocation, folding and trafficking in the ER and/or, (iii) activating genes that are involved in elimination of misfolded proinsulin through Endoplasmic Reticulum Associated Protein Degradation (ERAD). Studies demonstrate that UPR signaling is essential to preserve beta cell function and survival. If the UPR fails to restore proper proinsulin folding, beta cells undergo apoptosis. Defects in the biosynthetic network (for folding, processing, trafficking, and exocytosis) can also lead to fulminant beta cell failure (6; 7). Therefore, it is critical to understand the cellular components that promote human beta cell insulin homeostasis and how they are maintained in the presence of extracellular insults (8).

To date, transcriptomic, proteomic and metabolomic analyses attempting to catalog beta cell proteins that promote insulin synthesis, folding, trafficking and processing in whole human pancreatic islets have identified a large number of contributing metabolic and signaling pathway components (9-12). Furthermore, murine models have identified several ER resident proteins involved in proper proinsulin folding and processing, notably BiP, GRP170 and PDIA1 (5). Collectively, these studies indicate that proinsulin maturation is regulated by chaperones and catalyzed by ER oxidoreductases such as Ero1 (which generates stoichiometric amounts of H_2_O_2_ for every disulfide bond produced). Although these data provide insight into islet biology, the precise biosynthetic and folding machinery that directs proinsulin maturation in human islet beta cells still remains largely undefined.

Here, we employed an unbiased affinity purification mass spectrometry (AP-MS) approach to identify the central machinery responsible for proinsulin folding in human islet beta cells. The data reveal a rich proinsulin biosynthetic network that is remarkably conserved across a diverse group of donors. Among the proinsulin interactors identified were novel proteins, e.g. MYO18A; chaperones, e.g. GRP94 and BiP; co-chaperones, e.g. ERDJ5 and ERDJ3; as well as ER catalysts of disulfide bond formation, e.g. QSOX1 and PRDX4. Further investigation highlights a functional role for PRDX4 in defending proinsulin folding that may contribute importantly to protection against T2D. Thus, the AP-MS data provide a roadmap for functional dissection of the proinsulin biosynthetic network and its impact on beta-cell health.

## Results

### Definition of the Human Proinsulin Biosynthetic Interaction Network

To identify the physical interactions that dictate proper proinsulin synthesis and folding we first generated a series of monoclonal antibodies to human proinsulin (**Fig. S1A**) and chose a conformation specific monoclonal antibody that selectively recognizes human proinsulin by immunoprecipitation (IP) (20G11) in the presence of 1% Triton X-100, with negligible cross-reactivity to mature insulin (**Fig. S1B**). We recently showed that even healthy islets harbor a subset of proinsulin molecules that are misfolded, albeit at lower levels than for proinsulin mutants, which exhibit severe misfolding leading to Mutant *INS*-gene-induced Diabetes of Youth (MIDY) (13). Therefore, we tested whether 20G11 recognizes misfolded human proinsulin in addition to the properly folded proinsulin. COS1 cells were transfected with wild-type proinsulin or human proinsulin variants bearing MIDY point mutations that induce misfolding. 20G11 efficiently IP’ed all MIDY proinsulin mutants tested (**Fig. S1C**).

Proinsulin was affinity purified from human islets using 20G11 or control mouse IgG-coupled beads and subjected to Mass Spectrometry (AP-MS). Notably, IP of proinsulin provided beta-cell specificity in the context of intact islets, thereby avoiding the need for islet dispersal and beta cell purification methods that can stress the cells. Initially, two MS quantification methodologies were compared; Label Free (LF) and Tandem Mass Tag (TMT) isobaric labeling approaches (**Fig. S2**). Although numerous interactors were similarly identified with both technologies, the LF method outperformed TMT in terms of fold-change separation. The dynamic range of fold-change detected by LF was as high as c.a. 16 while the fold-change detected by TMT was compressed to 1 – 2.5, as described by others (14). Based upon optimal fold-change separation, we implemented LF analysis for subsequent studies.

With the goal of identifying highly conserved proinsulin interactions at the core of normal human beta cell function, we procured islet preparations from 6 donors that included Caucasian, Hispanic and African American ethnicities as well as both genders. The donors had no history of diabetes, BMI ranging from 21 to 25.4 and normal HbAIC (4.8%-5.5%) at the time of death (**Fig. 1A**). Equal numbers of islet equivalents were lysed and IP’ed with either 20G11 or mouse IgG conjugated beads, with replicates. Proinsulin IPs were subjected to on-bead denaturation, reduction and trypsin digestion prior to LC-MS/MS analyses.

**Figure 1.**
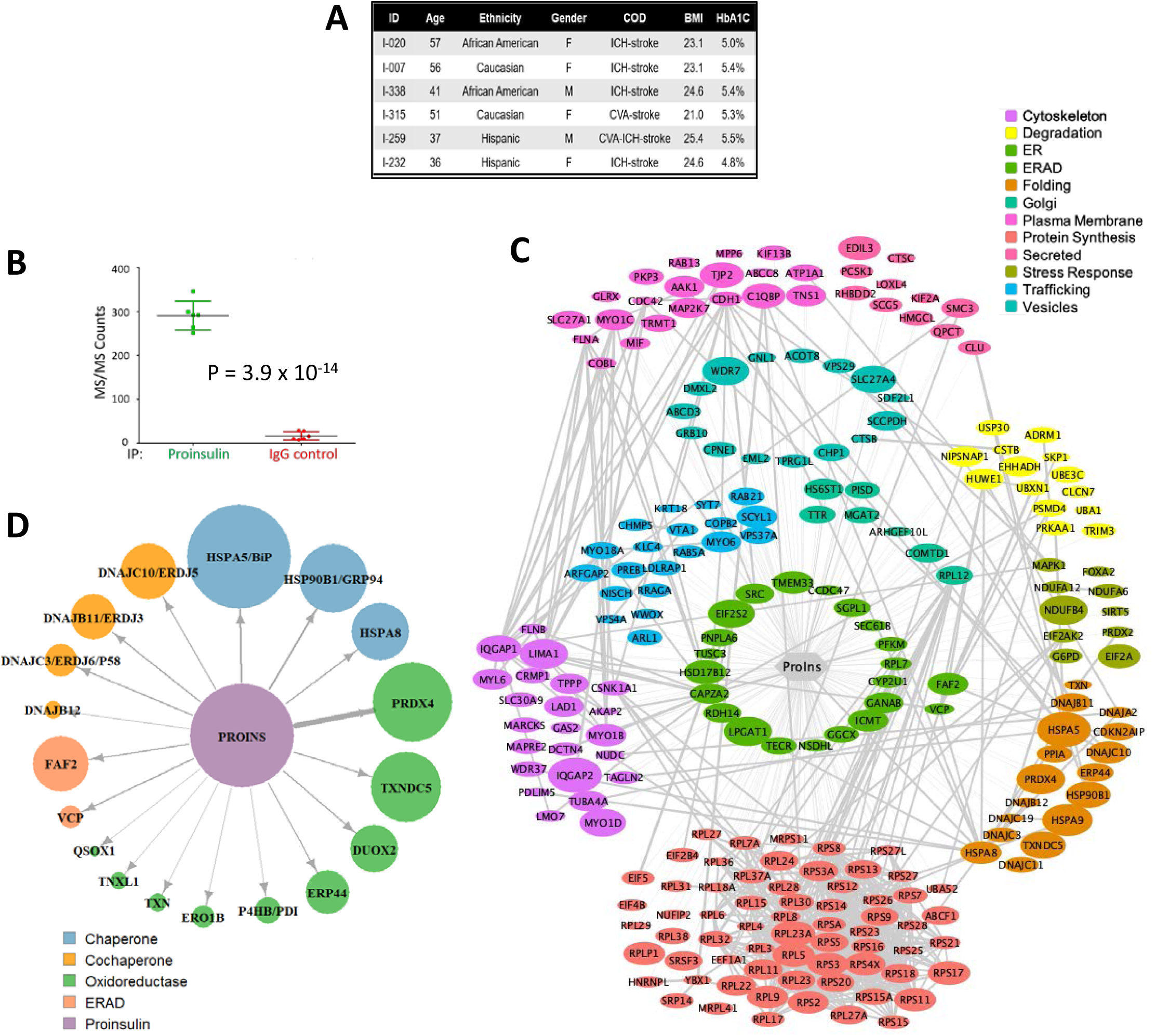
Defining the Proinsulin Biosynthetic Interaction Network. (A) Human islets from 6 non-diabetic donors were used for AP-MS. (B) Total MS/MS counts for Proinsulin bait (green) or IgG bait (red) from the six islet preparations reported in (A). (C) Network analysis of robust proinsulin protein:protein interactions generated by Cytoscape. Data were filtered for all of the following criteria (i) Proinsulin IP/Control IP intensity ratios ≥ 2fold, (ii) p ≤0.05, (iii) total MS/MS across 6 samples ≥10, and (iv) mRNA expression in single cell mRNA profiling of beta cells of at least 1 CPM (counts per million) (24). Colors indicate protein categories. The following categories; nuclear, mitochondria and undefined, are included in uploaded analysis but not in this figure. Increasing icon size depicts increased significance. (D) List of the most robust endoplasmic reticulum (ER) resident proinsulin interactors as determined by p value ≤0.05. Icon size reflects significance (smaller p values) and arrow thickness correlates with increasing log2FC Proinsulin IP/IgG IP.

MS/MS spectra were searched against the Human Uniprot database and mass spectra were analyzed with MaxQuant software (version 1.5.5.1; normalization and statistical analyses are described further in Methods). Signals were normalized using R’s Loess Pair Normalization. For statistical analyses, MSStats (15) was utilized to calculate a confidence score (*p*-value) for each protein based on the reproducibility of detection across samples. All data including *.raw data file, MaxQuant *.txt search results are available online at ProteomeXchange - Dataset ID PXD014476 (http://proteomecentral.proteomexchange.org/cgi/GetDataset). Comparison of MS/MS counts for bait (proinsulin) in proinsulin IP versus control IP values across the 6 individual samples revealed remarkable consistency in the recovery of proinsulin, suggesting little technical variability (**Fig. 1B**).

To identify the most robust proinsulin interactions for network analysis, the data were stringently filtered using the following criteria; (i) Proinsulin IP/Control IP intensity ratios ≥ 2-fold, (ii) *p* ≤ 0.05, and (iii) total MS/MS across six samples ≥ 10. Moreover, to be certain that identified interactors were derived from beta cells, proteins were removed from analysis if their mRNA expression, in a recent beta cell single cell mRNA profiling study, did not reach a minimum threshold of 1 CPM (counts per million) average across all beta cell samples (16). The resulting dataset identified 461 proteins from multiple subcellular compartments, whose functions include protein synthesis, folding, degradation, trafficking, secretory vesicle formation, and proteolytic cleavage to mature insulin, as well as proteins of unknown function (**Fig. 1C**). For visualization of the network, cytosolic, nuclear, and mitochondrial proteins were removed as described in reference (17), although they remain in the uploaded dataset, which can be found at ProteomeXchange (See above). Thus, we constructed the first human proinsulin biosynthetic network (**Fig. 1C**).

Among the identified proinsulin interactors was the proteolytic enzyme, PCSK1 (Prohormone Convertase 1), that cleaves the C-peptide/B chain junction of proinsulin. Identifying the proinsulin: PCSK1 association with high confidence (MSStat *p* value = 4.6 x 10^−5^, log_2_FC = 1.47), provides a positive control validating the confidence in the sensitivity of the interaction data. Conversely, Islet Amyloid Polypeptide (IAPP), a highly expressed beta-cell protein that is stored and co-secreted with insulin but is not thought to physically interact with proinsulin, was not identified among the prey proteins, which speaks to the specificity of the affinity purification.

Proinsulin immunoprecipitations and Western blots were used to validate select interactors **(Fig. S3**). Among the novel interactors was the Unconventional Myosin-XVIIIa (MYO18A, MSStats *p* value=5.86×10^−6^, log_2_FC = 3.71, 98.9^th^ percentile) (**Fig. S3B**). To the best of our knowledge, MYO18A has not been studied in beta cells. MYO18A is thought to link Golgi membranes to the cytoskeleton, participating in the tensile force required for vesicle budding from the Golgi (18). Therefore, our finding that MYO18 interacts with proinsulin, even if indirectly, supports the idea that there is specific interaction (that withstands 1% Triton X-100) and recruitment of proinsulin at the luminal aspect of budding Golgi membranes, as previously suggested (19).

### The ER localized proinsulin interaction network

We recently showed that a subpopulation of proinsulin in normal human islets is misfolded within the ER, existing as disulfide-linked oligomers and higher molecular weight complexes in addition to properly folded proinsulin (3). Further, we found that proinsulin misfolding increased during progression to diabetes, and was the first alteration detected in the beta cell prior to altered glucose homeostasis (3). Therefore, we focused on proinsulin interactors that may regulate proinsulin folding in the ER. In order to capture ER resident proteins that are likely to interact transiently (or only with a misfolded subset of the larger population of proinsulin molecules), we relaxed stringency [using ß-cell expression criteria as described above and p ≤ 0.05 (**Fig. 1D**)].

Not surprisingly, the well-known chaperone BiP (HspA5), thought to be required for productive proinsulin folding (20), was identified as one of the most statistically significant ER protein interactors with proinsulin (**Fig. 1C and D**). Direct interaction of BiP with proinsulin was previously shown to increase with increased proinsulin synthesis (21). In addition to BiP, we also found interactions between the ER chaperones GRP94 and co-chaperones ERDJ3 and ERDJ5 and proinsulin (**Fig. 1C and D**), confirmed by AP-western (**Fig. S3A and B**).

Recent studies in model organisms suggest that multiple ER enzymes may catalyze protein disulfide bond formation (4; 22), however the expression and role of these enzymes in human ß-cells remains unexplored. Here, AP-MS revealed that human proinsulin associated with QSOX1, validated in **Fig. S3D**, an enzyme with no previously known substrates (23). Proinsulin also interacted with Peroxiredoxin-4 (PRDX4) (**Figs. 1C and D)**. PRDX4 is a 2-cysteine peroxiredoxin that utilizes luminal H_2_O_2_ to oxidize proteins (17). In fact, PRDX4:proinsulin interactions were among the most robust in our MS dataset (p=4.92×10^−11^, Log_2_FC = 6.17, 99.6^th^ percentile) (**Fig. 1D**). Expression of PRDX4 in human beta cells was confirmed in a recent single cell mRNAseq study of human islets (24) (**Fig. S4**), enabling further investigation into its role in proinsulin folding.

### Functional characterization of PRDX4 suggests a role in supporting proper proinsulin folding

To validate the proinsulin:PRDX4 interaction, HEK293A cells were co-transfected with Flag-tagged PRDX4 and MYC-tagged wild-type or *Akita* proinsulin. Proinsulin-MYC pull-down co-precipitated PRDX4 and conversely, PRDX4 pull-down co-IP’ed a major fraction of proinsulin. Moreover, PRDX4 efficiently precipitated MYC-tagged *Akita* mutant proinsulin, revealing that PRDX4 interacts with both folded and misfolded proinsulin (**Fig. S3C**).

Given the major role of BiP in the beta cell’s ER (21), we considered that PRDX4 might be a component of a BiP containing complex. AP-Westerns revealed that PRDX4 indeed bound to BiP (**Fig 2A**), although the interaction did not require proinsulin. We next asked whether BiP was required for the proinsulin:PRDX4 interaction. For this experiment we treated cells with the BiP ATPase inhibitor HA15, or the subtilase cytotoxin SubAB that cleaves and inactivates BiP (25). As a negative control, we employed mSubAB, an enzymatically inactive form of SubAB (**Fig 2B**). PRDX4 interactions were unchanged by BiP inhibition or cleavage, evidence that intact or enzymatically active BiP is not required for PRDX4:proinsulin interactions (**Fig 2B**).

**Figure 2.**
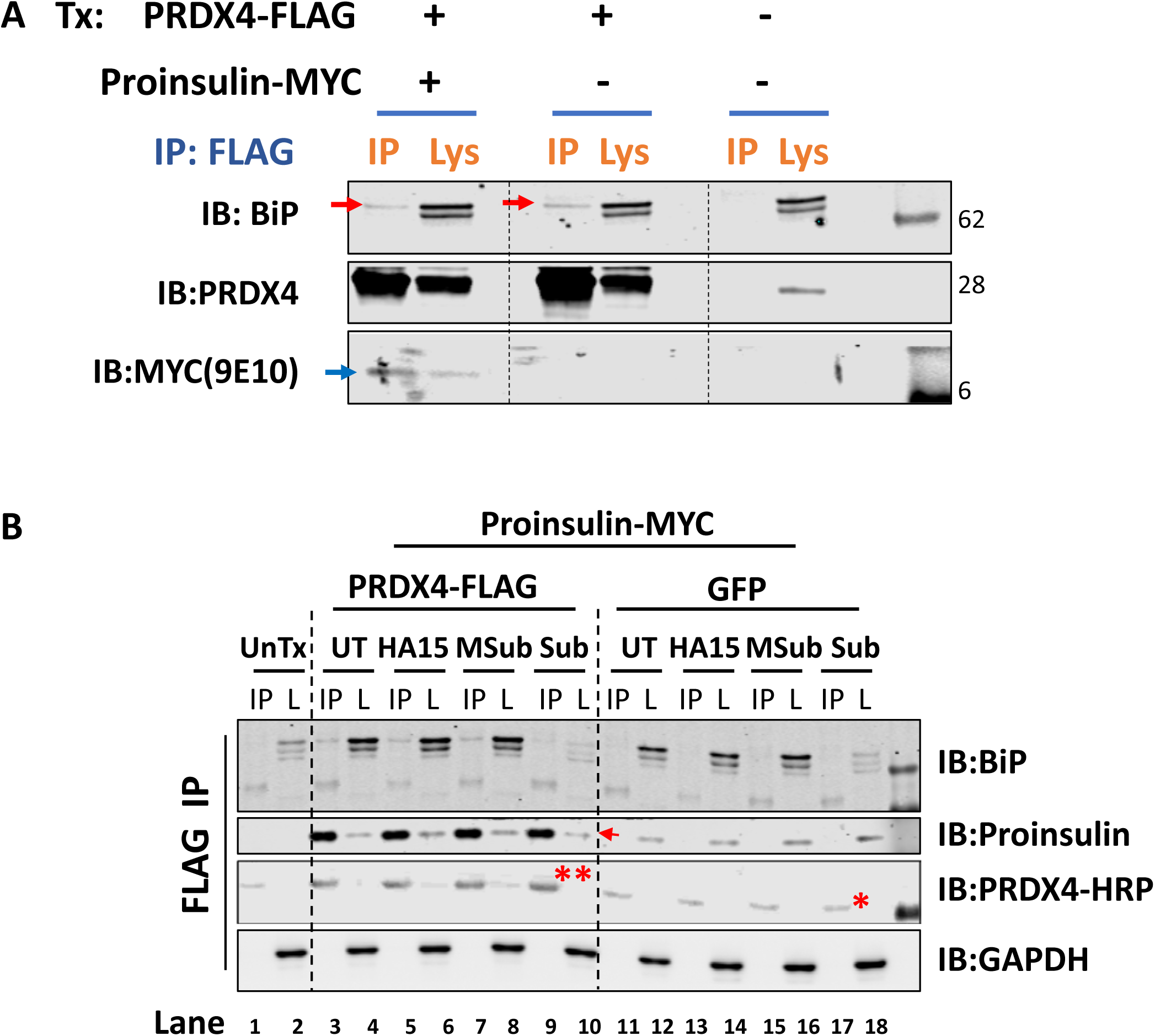
PRDX4 interactions with BiP and Proinsulin are independent of each other. (A) HEK293A cells were transfected with combinations of control pCDNA (-), Proinsulin-MYC and PRDX4-FLAG. Resulting lysates were subject to immunoprecipitation with anti-FLAG magnetic beads. Lysates and IP fractions were immunoblotted for BiP, PRDX4 or MYC (proinsulin). Red arrows show BiP and blue arrow identifies Proinsulin-MYC. (B) HEK293T cells were transfected with combinations of Proinsulin-MYC, PRDX4-FLAG and GFP. After 44 hours cells were treated with SubAb (2 µg/mL, 4 hrs), Mutant SubAb (2 µg/mL, 4 hrs), HA15 (10 µM, 4 hrs) or untreated, followed by immunoprecipation with anti-FLAG magnetic beads. 3% of Lysate (L) and 20% of immunoprecipitate (IP) were analysed on reducing SDS- PAGE gels. Arrow identifies proinsulin band IP’d with PRDX4-FLAG, (*) is IgG band in PRDX4 lanes and (**) is PRDX4 specific band, appearing at the same molecular weight as IgG but with higher intensity.

To investigate the conformational composition of PRDX4 in islets, we analyzed murine islets by nonreducing SDS-PAGE and immunoblotting for PRDX4. Virtually all PRDX4 protein was found in disulfide linked dimers at approximately 52 KDa and higher molecular weight complexes (HMW), at 100 KDa, with only a small fraction as the 28 KDa monomer (**Fig. S5A**). To validate the specificity of the HMW PRDX4 complexes, the islets were treated with increasing concentrations of dithiothreitol (DTT, a reducing agent) *in vivo* (**Fig. S5A**). High DTT conditions resolved PRDX4 HMW species to monomer, while retaining a particularly stable dimer band at 52kDa **(Fig. S5A)** (22). Because PRDX4 was previously identified in a PDI trap assay we considered that its ability to enter into the observed HMW complexes might depend on PDIA1. We therefore repeated the study in islets from mice with beta cell specific deletion of PDIA1, finding that PRDX4 complex formation was not altered by PDIA1 loss and that loss of PDIA1 actually increased PRDX4 expression (**Fig. S5 B**).

To determine whether proinsulin folding is assisted by PRDX4, lentiviral vectors expressing shRNA were used to deplete PRDX4 in MIN6 cells, achieving 64 – 75% PRDX4 knockdown compared to controls (**Fig. 3A and B**). PRDX4 knockdown triggered proinsulin misfolding as revealed by an increase of high molecular weight complexes and moreover, PRDX4-depleted ß-cells were hypersensitive to proinsulin misfolding induced by oxidant challenge with 100 µM menadione or 1 mM H2O2 for 1 hour **(Fig. 3A**). Decreased PRDX4 expression also resulted in increased mRNA levels of CHOP, a transcription factor associated with UPR, suggesting that loss of PRDX4 elevated ER stress (**Fig. 3C**). We did not however, observe a change in cell proliferation or cell survival (not shown).

**Figure 3.**
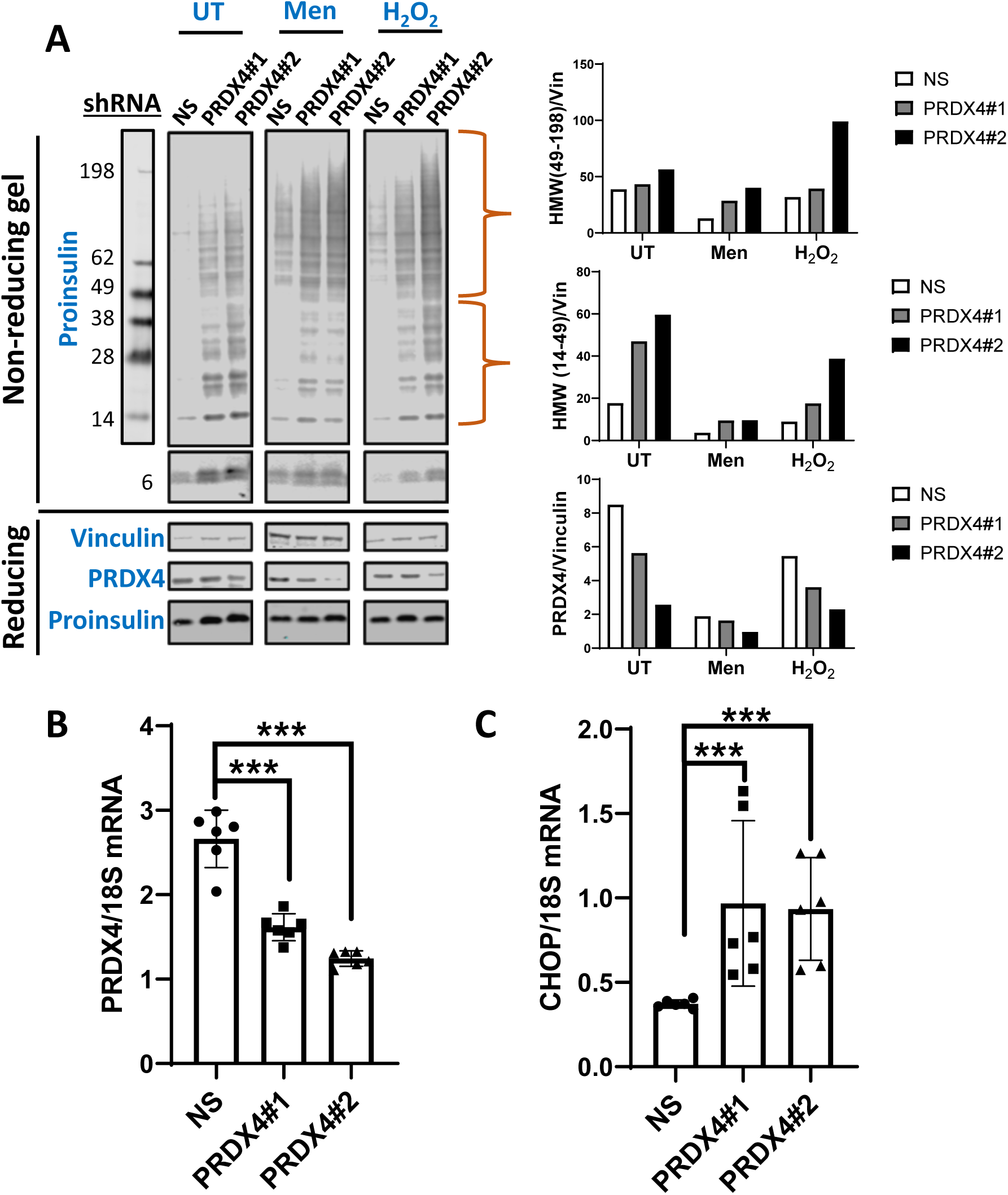
Loss of PRDX4 exacerbates proinsulin misfolding induced by oxidative stress in MIN6 cells. MIN6 cells stably expressing PRDX4 shRNA (PRDX4#1 and PRDX4#2), or scrambled non-specific shRNA (NS) were generated. (A) Cells were untreated with menadione (Men) (100 µM, 1 hr) or H_2_O_2_ (1 mM, 1 hr). Non-reducing gels were blotted with proinsulin antibody CCI17. Bottom panels of non-reducing gels shows darker exposure of non-reducing proinsulin monomer. Reducing gels were blotted for Vinculin, total Proinsulin, and PRDX4. Accompanying graphs (upper right) showed quantification of HMW bands from 14 – 49 KDa, 49 – 198 KDa and total PRDX4 normalized by Vinculin. (B) PRDX4 knockdown is confirmed by qPCR using 3 biological replicates per line with technical duplicates for each biological replicate to ensure reproducibility. (C) Analysis of the ER stress marker CHOP by qPCR.

We next asked whether increased expression of PRDX4 could promote proper proinsulin folding. Here HEK293T cells were co-transfected with MYC-tagged proinsulin and either FLAG-tagged PRDX4 or GFP control (**Fig. 4**). Western blots revealed that exogenous PRDX4 expression increased the ratio of properly folded proinsulin monomer and low molecular weight oligomers (<50kDa) versus high molecular weight species (>98 KDa) (**Fig. 4**).

**Figure 4.**
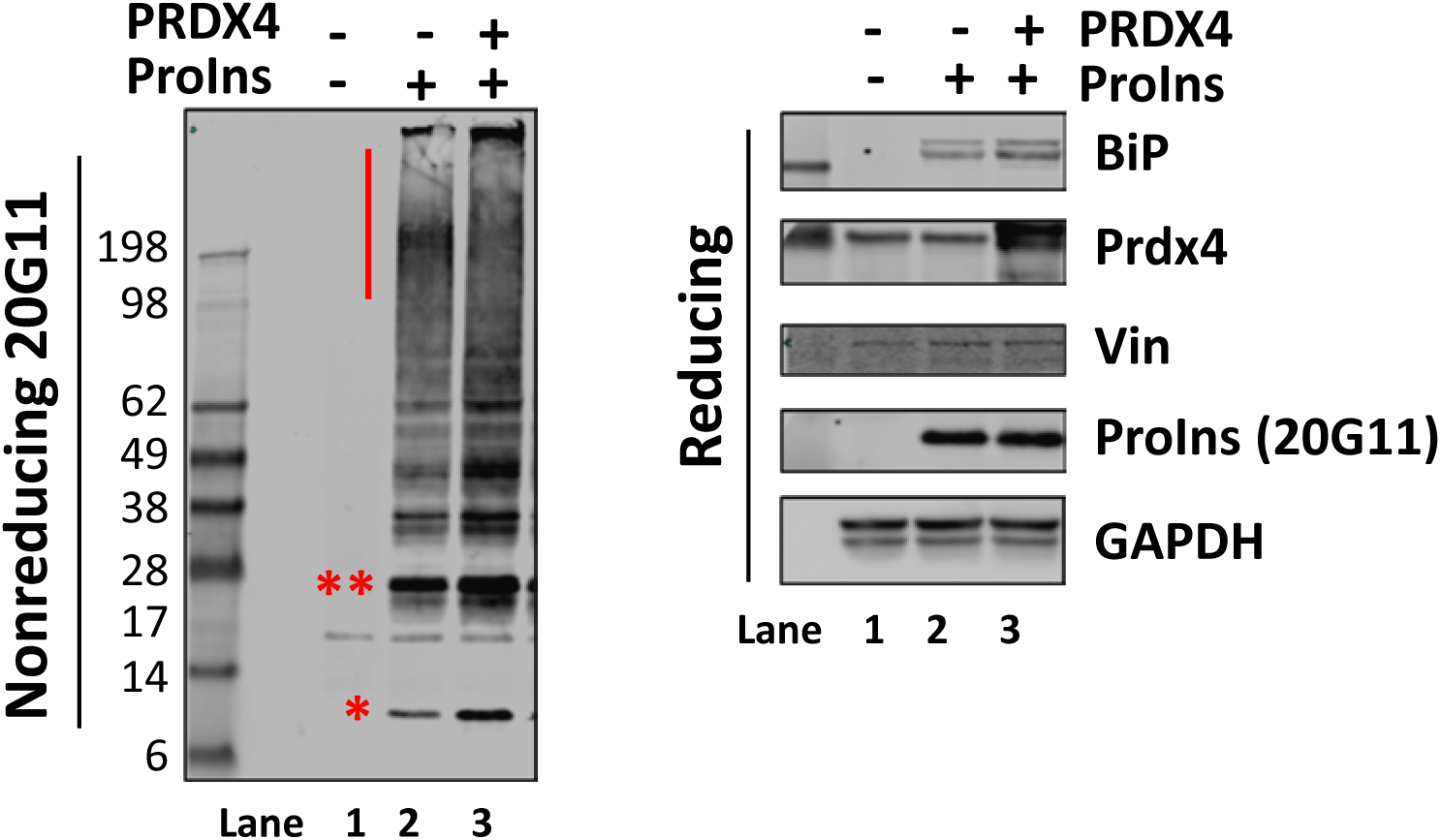
Exogenous PRDX4 promotes proper proinsulin folding. HEK293T cells were transfected with combinations of Proinsulin–MYC, PRDX4-FLAG and GFP for 48 hours prior to harvesting cells and SDS-PAGE Analysis. On the left is non-reducing gel, red bar indicates high molecular weight complexes (>98 KDa). (*) and (**) denote monomer and dimer proinsulin bands, respectively. On the right is the accompanying reducing gel immunoblotted for BiP, PRDX4, total Proinsulin and loading controls (Vinculin, GAPDH). The experiment was repeated 3x.

Given the increased level of proinsulin misfolding caused by external insults in HEK and MIN6 cells, we investigated the effects of Menadione, an oxidant (4) or, Brefeldin A (BFA) (26; 27) on conformational changes in proinsulin and PRDX4 in the context of human islets (**Fig. S6**). Treatment of human islets with increasing concentrations of either Menadione or BFA induced proinsulin misfolding compared to untreated human islets (**Fig. S6**). Only oxidative stress, however recruited PRDX4 into high molecular weight disulfide bonded complexes (**Fig. S6**) suggesting that PRDX4 is recruited to client proteins during oxidative stress.

The expression of multiple ER proteins increases in response to glucose stimulation of beta cells (28). We therefore asked whether PRDX4 expression in human islets was glucose responsive. Islets were starved in glucose free media for 1 hour before treatment with low (3 mM) or high (33 mM) glucose for 18 hours. As expected, high glucose stimulated insulin release into the media (**Fig. 5A**). The observed GSIS corresponded to a 45% increase in the steady state level of proinsulin; and a 2.3x increase in BiP (**Fig. 5C**), a chaperone with known glucose responsive expression (29). In contrast, PRDX4 expression remained constant across all samples (**Fig. 5C**). Notably, non-reducing gels showed that the upregulated proinsulin biosynthesis led to an increase in the ratio of HMW (misfolded) proinsulin relative to monomer in human islets (**Fig. 5A**).

**Figure 5.**
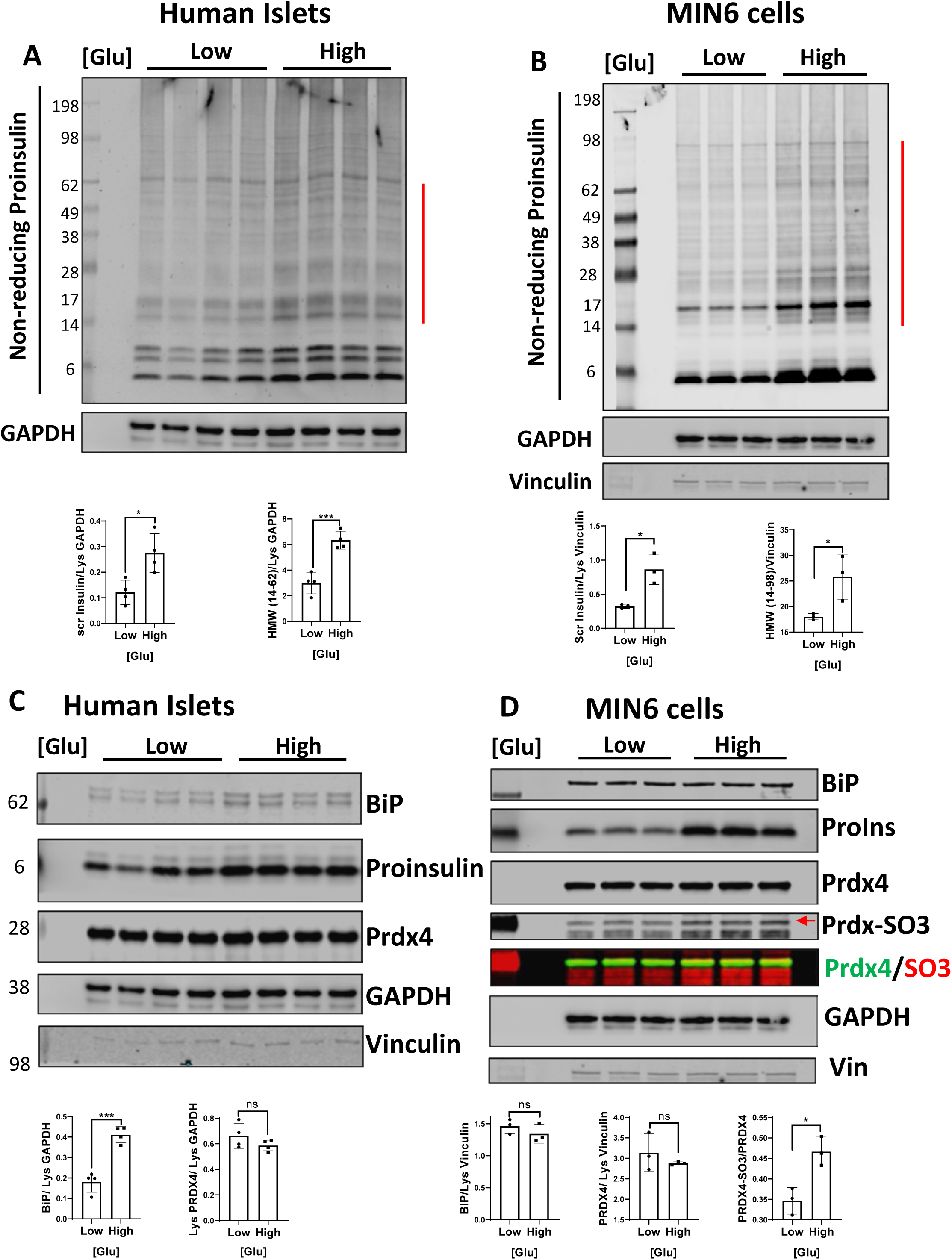
High glucose induces proinsulin misfolding in human islets and MIN6. (A) High glucose (High = 30 mM vs. Low = 3 mM) induces not only secreted insulin but also high molecular weight proinsulin complexes (indicated by vertical red bar). Accompanying graphs show secreted insulin in the media and high molecular weight complexes, normalized by GAPDH in the lysate, respectively. (B) High glucose (High = 22 mM vs. Low = 2.2 mM) induces HMW complexes (red bar) in MIN6 cells. Accompanying graphs show secreted insulin in the media and high molecular weight proinsulin complexes, normalized by Vinculin in the lysate, respectively. (C) Samples described in (A) were analyzed on reducing SDS-PAGE and immunoblotted for BiP, total proinsulin, PRDX4 along with loading controls (Vinculin and GAPDH). Accompanying graphs show the levels of BiP and PRDX4 normalized by GAPDH. (D) Samples described in (B) were analyzed on reducing SDS-PAGE and immunoblotted for BiP, total proinsulin, PRDX4, and PRDX-SO_3_ along with loading controls (Vinculin and GAPDH). PRDX4 and PRDX-SO3 were immunoblotted with two different secondary antibodies: green = PRDX4 and red = PRDX-SO_3_ on the same membrane and merged. Arrow identifies PRDX4-SO_3_ band. Accompanying graphs indicate BiP and PRDX4 levels normalized by GAPDH, and PRDX4-SO_3_ normalized to total PRDX4.

The experiment was similarly performed in MIN6 cells with low (2.2 mM) versus high (22mM) glucose for 22.5 hours (**Fig. 5B**). High glucose treatment in MIN6 cells resulted in a 3.3 fold increase in secreted insulin accompanying a 22% increase in intracellular steady state levels of proinsulin (**Fig. 5B**). As in human islets, elevated glucose also induced modest proinsulin misfolding, and PRDX4 expression was not altered by glucose stimulation (**Fig. 5D**). Moreover, in MIN6 cells, the high glucose-induced proinsulin misfolding was accompanied by sulfonylation of PRDX4 (**Fig. 5D**), a modification known to inactivate the catalytic activity of PRDX4 (30). Intriguingly, subsequent examination of PRDX4 sulfonylation status in a series of untreated human islet samples from diabetic (n=4) and non-diabetic (n=4) sources revealed for the first time that islets from patients with T2D exhibit 3.3-fold increase in sulfonylated PRDX4 relative to total PRDX4 (p=0.011) (**Fig 6**).

**Figure 6.**
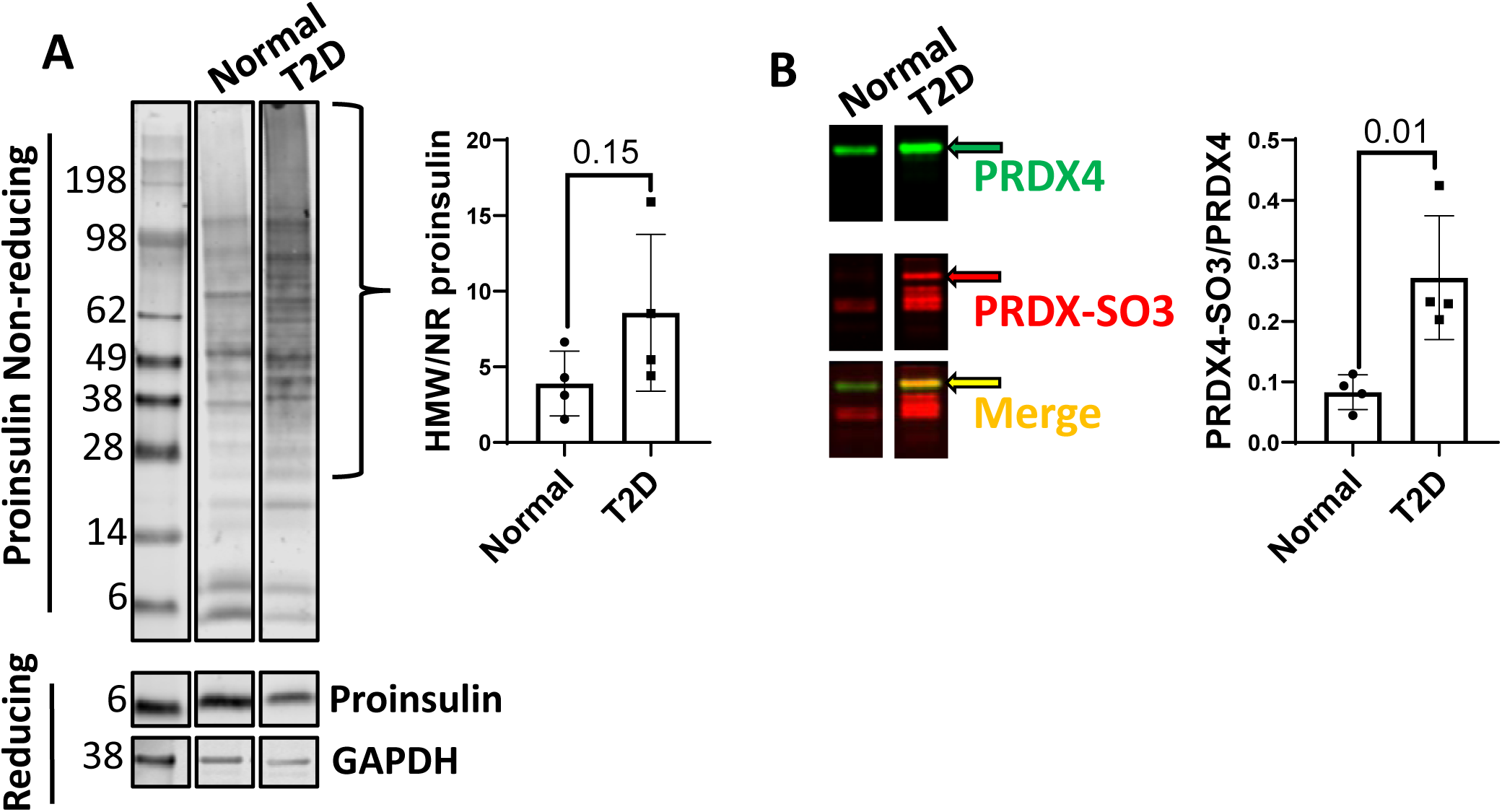
T2D islets trend to increased proinsulin misfolding and exhibit increased PRDX4 sulfonylation. (A) Non-reducing gels (upper panel) of human islet preparations from 4 individual normal and 4 Type 2 Diabetic (T2D) islet preparations were run, normalized by total protein concentration determined by Lowry assay, and blotted for human proinsulin (all four samples were analyzed, one representative sample is displayed here). Reducing gels immunoblotted for total proinsulin and loading control (GAPDH) are shown (bottom panels). The accompanying graph shows quantification of HMW bands (size range indicated by curved bracket) normalized by proinsulin monomer on the non-reducing gel for all 8 samples. (B) Samples analyzed in (A) were immunoblotted for both PRDX4 (green) and sulfonylated PRDXs (i.e. PRDX-SO_3_, red). Merged image showed co-localization of PRDX4 band with its sulfonylated version at the same molecular weight. Intensity of the sulfonylated PRDX4 bands (yellow arrow) versus total PRDX4 were quantified.

## Discussion

Significant efforts have been made to understand beta cell biology at OMICs level. One potential limitation of these studies is that they may inadvertently incorporate results from multiple cell types in the pancreas despite efforts to purify beta cells. Advances in OMICs at the single cell level are playing an important role in defining cells at the RNA level but the single cell approach remains difficult with proteins. By focusing specifically on the proinsulin folding/processing pathway, we were able to generate beta-cell specific data in the context of whole islets and thus avoided harsh conditions to disperse islets, which may alter beta cell physiology. To focus on synthesis and trafficking, we generated a highly specific monoclonal antibody to human proinsulin, that efficiently immunoprecipitates proinsulin with negligible reactivity to mature insulin (**Fig S1**).

The most striking feature of this data set is the tight conservation of human proinsulin biosynthetic network across 6 donors reflecting 3 ethnicities and both genders. A technical aspect that may have contributed to the data concordance was that human islets were procured from a single source to avoid artifacts due site-specific islet isolation practices.

For the network design the criteria were quite stringent, requiring (i) at least 2 fold increase in Intensity between 20G11 IP and IgG IPs, (ii) p ≤ 0.05 and, (iii) MS/MS counts of at least 10 among the 6 human islet preparations, as well as (iv) a minimum mRNA expression level in published mRNA seq studies from single beta cells (24). Our findings corroborate the previously identified proinsulin interacting proteins, BiP (HSPA5) and GRP94 (GSP90B1) that have both been shown to play essential roles in proinsulin folding (21; 31) while extending those studies to provide an entire interaction network.

We previously performed AP-MS to identify proinsulin and insulin interactors in murine MIN6 cells (32). Consistent with those studies, here we identified DNAJB11(ERDJ3), DNAJC6 (p58^IPK^), and ERP44 as high confidence human proinsulin interacting proteins. Not surprisingly, TMEM24, that interacted more robustly with mature insulin than with proinsulin in our previous murine MIN6 study, was not identified in this study, which focused solely on proinsulin vs insulin interactions in human islets. The two studies also utilized unique, conformation-specific, antibodies, which may lead to differences in prey captured.

Among the most significant newly identified proinsulin interacting proteins was PRDX4, the only one of the 6 human PRDXs that is ER resident. We show in native islets that PRDX4 primarily resides in disulfide linked dimers and high molecular weight complexes which may include oligomers (22) but also clearly involve other proteins as not all bands are multimers of 28 KDa. Importantly, all bands collapse to PRDX4 monomer and dimer upon treatment with DTT *in vivo* in murine islets, validating that they specifically contain PRDX4.

Although PRDX4 is not an essential ER enzyme in mice, except in testes (33), we find that proinsulin is more prone to misfolding when PRDX4 levels are low and stress is induced by oxidant treatment (**Fig. 3**), indicating that under stress conditions PRDX4 plays an important role. These data are consistent with the finding that MEFs lacking ERO1 were intolerant of PRDX4 deficiency (17). The involvement of PRDX4 with the redox protein network is further emphasized by the finding that loss of QSOX1, one of our identified proinsulin interactors induces overexpression of PRDX4 in mouse hearts (34). Here we also show that loss of PDIA1 increases expression of PRDX4 suggesting that there is increased need for PRDX4 in the absence of PDIA1. Whether islets from PRDX4 knockout mice would exhibit proinsulin misfolding under stress is an avenue for investigation.

In yeast, it was suggested that the single PRDX-like protein is a molecular triage agent that absorbs oxidation to spare thioredoxin (35). It may not be surprising then that overexpression of PRDX4 in mice repressed beta-cell apoptosis, induced beta-cell proliferation and provided protection to beta-cells under stress, e.g. against streptozotocin-induced diabetes (36).

Excessive oxidation was shown to inactivate PRDX4 by sulfonylation (30). In MIN6 cells, high glucose alone induces sulfonylation of a portion of PRDX4 (**Fig. 5**). Notably, we show that sulfonylation of PRDX4 is increased in islets from patients with T2D (**Fig. 6**). The data suggest that beta cells in patients with T2D have increased oxidative stress and/or diminished capacity to handle oxidative stress, which is in agreement with previous studies (37; 38). It is intriguing that serum of patients with T2D have higher levels of circulating PRDX4 than controls (39). It will be of interest to determine whether circulating PRDX4 emanates from beta cells and/or whether (13; 40) it is modified by sulfonylation.

In summary, we have identified a complex set of protein-protein interactions that reflects the dynamic folding compartments that dictate proinsulin folding and trafficking through the exocytic pathway (41; 42) in human beta cells. The proinsulin interaction network provides a critical resource to characterize previously unknown molecular features of the beta cell secretory pathway. It will now be of interest to determine which proteins interact directly versus indirectly with proinsulin and to uncover their individual roles in promoting efficient proinsulin biosynthesis.

## Methods

### Cell Culture and Treatments

Islets were procured from Prodo labs (Aliso Viejo, CA). Islet preparations averaged 85-95% purity by DTZ and 95% viability by EB/FDA staining. Each preparation was tested for glucose responsiveness prior to use. Islets were cultured in Prodo Islet Complete Media, Prodo labs (Aliso Viejo, CA). MIN6 cells were cultured in DMEM with 4.5g/L glucose and L-glutamine, 3.4% NaHCO_3_, 1X penicillin, streptomycin, 275 nM β-mercaptoethanol (BME), 15% FBS. HEK 293A and 293T cells were cultured in DMEM with 4.5g/L glucose and L-glutamine, sodium pyruvate, 1X penicillin, streptomycin, 10% FBS. Perturbagent conditions used were menadione (100 µM, 1 hr), H_2_O_2_ (1 mM, 1 hr), Brefeldin A (5 µg/mL, 1 hr) or SubAB/Mutant SubAB (2 µ/mL, 4 hrs), HA15 (10µM, 1hr). After treatment, cells were washed once with PBS. Islets and MIN6 cells utilized for AP-western were typically lysed in RIPA or Lysis Buffer as described below. MIN6 and HEK cells utilized for SDS-PAGE analysis were lysed directly in 2X Laemmli without BME, heated at 100°C for 10 mins and clarified by centrifugation at 17,000 g for 15 minutes. For reducing samples, BME was added to lysate aliquots to 2.5% and heated again at 100°C for 10 mins before SDS-PAGE.

### Immunoprecipitation of Proinsulin from Human Islets for mass spectrometry analyses

Human islets were lysed in lysis buffer (50mM Tris pH7.4, 150mM NaCl and 1% TX-100, and 1X protease inhibitor cocktail (Thermo Scientific Pierce)) on ice for 45 min. For each IP used in MS, lysates were pre-cleared with Protein G beads. Lysate was immunoprecipitated with beads crosslinked to mouse IgG or proinsulin antibody (20G11) overnight at 4°C and then washed twice with lysis buffer and once with lysis buffer without detergent. A fraction of the beads was removed for protein elution to confirm successful IP by western blot and silver staining. The majority of the beads were subjected to denaturation, reduction and trypsin digestion followed by Mass spectrometry analysis.

### Mass spectrometry sample preparation

Proteins were digested in 8M urea 50 mM ammonium bicarbonate buffer. Briefly, cysteine disulfide bonds were reduced with 5 mM tris(2-carboxyethyl)phosphine (TCEP) at 30°C for 60 min followed by cysteine alkylation with 15 mM iodoacetamide (IAA) in the dark at room temperature for 30 min. Following alkylation, urea was diluted to 1 M using 50 mM ammonium bicarbonate, and proteins were finally subjected to overnight digestion with mass spec grade Trypsin/Lys-C mix (Promega, Madison, WI). Digested proteins were finally desalted using a C18 TopTip (PolyLC, Columbia, MD) according to the manufacturer’s recommendation), and the organic solvent was removed in a SpeedVac concentrator prior to LC-MS/MS analysis. To label samples with TMT for TMT vs. LF comparison, TMTsixplex™ Isobaric Label kit (ThermoFisher Scientific) was used according to manufacture protocol.

### Mass spectrometry data acquisition

For 3 out of 6 islet batches, islets were divided into four equal parts based on number of islet equivalents, one of which was immunoprecipitated with mouse IgG as a control, and the other three with the proinsulin specific antibody 20G11 as three experimental replicates. Prior to MS/MS analyses, three experimental replicates of 20G11 IP were pooled together resulting in a total of 2 samples. Each sample was subject to state of the art 2 Dimensional LC-MS/MS analysis in which the peptide mixture was first separated in a high-pH reverse phase chromatography into 5 fractions before directly subject to a second low-pH reverse phase chromatography coupled with MS/MS analyses. This analysis results in 5 technical replicates per sample and a total of 30 LC-MS/MS runs. For the other 3 islet batches, islets were also divided into four equal parts, two of which was precipitated with IgG as a control and the other with 20G11. Each experimental replicate was analyzed separately in LC-MS/MS analysis resulting in a total of 60 runs with 10 technical replicates per sample.

Dried samples were reconstituted in 100mM ammonium formate pH ∼10 and analyzed by 2DLC-MS/MS using a 2D nanoACQUITY Ultra Performance Liquid Chromatography (UPLC) system (Waters corp., Milford, MA) coupled to an Orbitrap Velos Pro mass spectrometer (Thermo Fisher Scientific). Peptides were loaded onto the first dimension column, XBridge BEH130 C18 NanoEase (300 µm × 50 mm, 5 µm) equilibrated with solvent A (20mM ammonium formate pH 10, first dimension pump) at 2 µL/min. The first fraction was eluted from the first dimension column at 17% of solvent B (100% acetonitrile) for 4 min and transferred to the second dimension Symmetry C18 trap column 0.180 × 20 mm (Waters corp., Milford, MA) using a 1:10 dilution with 99.9% second dimensional pump solvent A (0.1% formic acid in water) at 20 µL/min. Peptides were then eluted from the trap column and resolved on the analytical C18 BEH130 PicoChip column 0.075 × 100 mm, 1.7µm particles (NewObjective, MA) at low pH by increasing the composition of solvent B (100% acetonitrile) from 2 to 26% over 94 min at 400 nL/min. Subsequent fractions were carried with increasing concentrations of solvent B. The following 4 first dimension fractions were eluted at 19.5, 22, 26, and 65% solvent B. The mass spectrometer was operated in positive data-dependent acquisition mode. MS1 spectra were measured with a resolution of 60,000, an AGC target of 10^6^ and a mass range from 350 to 1400 m/z. Up to 5 MS2 spectra per duty cycle were triggered, fragmented by CID, with an AGC target of 10^4^, an isolation window of 2.0 m/z and a normalized collision energy of 35. Dynamic exclusion was enabled with duration of 20 sec.

### Mass Spectrometry Data Processing

All mass spectra from were analyzed with MaxQuant software version 1.5.5.1. MS/MS spectra were searched against the Homo sapiens Uniprot protein sequence database (version January 2015) and GPM cRAP sequences (commonly known protein contaminants). Precursor mass tolerance was set to 20ppm and 4.5ppm for the first search where initial mass recalibration was completed and for the main search, respectively. Product ions were searched with a mass tolerance 0.5 Da. The maximum precursor ion charge state used for searching was 7. Carbamidomethylation of cysteines was searched as a fixed modification, while oxidation of methionines and acetylation of protein N-terminal were searched as variable modifications. Enzyme was set to trypsin in a specific mode and a maximum of two missed cleavages was allowed for searching. The target-decoy-based false discovery rate (FDR) filter for spectrum and protein identification was set to 1%. Note that proinsulin is searched as insulin in the database.

### Normalization

Intensity of each identified peptide was subtracted from the control IP and Log2Fold Change Proinsulin IP/IgG IP (Log_2_FC) was calculated. Two different methods of Mass Spectrometry analysis, Label Free and Tandem Mass Tag (TMT) isobaric labeling analysis were employed for a single islet prep (HP 15157-01) to compare the two methods (**Figure S2**). Based on resolution of fold changes, Label Free (LF) analyses were implemented for all of the samples.

For normalization, Loess Global vs Pair, quantile, MAD and Total Intensity methods were compared. Preliminary evaluation of the most suitable normalization method for the IP-MS data (Loess-R) was conducted with Normalyzer (43). Although outcomes looked very similar with all of these methods Loess Pair gave the best normalized dataset as the method considers control and sample within each experiment separately rather than averaging samples and controls across subjects. This is most suited for human islets given the possibility of variations from one human subject to another.

### Statistical Analysis

For Statistical analysis, the datasets were analyzed by MSStats (15) to score and evaluate confidence of interactions in proinsulin IPs versus the control IgG coated beads. MSStats calculates Fold change and allows one to extract a confidence score (p-value) for each protein based on the reproducibility of detection across samples. Due to a large dynamic range in the demographics of our islet donors, MSStats was chosen as this open source R based platform tool enables better quantitation of the sources of variations (44).

Final list of proinsulin interactome was selected to have (i) Fold Change (Ins vs. Control) ≥2, (ii) p-value ≤ 0.05 and (iii) deltaMS/MS (total sum of spectral counts across replicates) ≥ 10. Additional filtering was performed using previously published single cell RNA-seq data (24) to generate list of genes that were “at least expressed in Beta-cells”. The threshold was set to 1 CPM (counts per million) average across all beta cell samples yielding ∼13K genes out of 20K.

### Bioinformatic Analyses

Initial proteostasis compartment analysis was comprised of unbiased and curated data as shown at https://github.com/balchlab/PPT_annotation. Pathway and process enrichment analysis were carried out with the following ontology sources: KEGG Pathway, GO Biological Processes, Reactome Gene Sets and CORUM. Gene annotation and analysis resources included Metascape (45) and Enrichr (46). Protein Atlas (47) was used as primary source for subcellular localization, protein function and proteostasis classification.

### Mice and Murine Islet Isolation

Mouse models used in this experiment were made as reported elsewhere (4). In brief, mice with PDIA1 floxed alleles were obtained from Dr. J. Cho (Univ. of Illinois-Chicago) and crossed with Rat Insulin Promoter (RIP)-CreER transgenic mice. PDIA1 deletion was performed by IP injection of the estrogen receptor antagonist Tamoxifen (Tam) (4mg/mouse) three times a week. All procedures were performed by protocols and guidelines reviewed and approved by the Institutional Animal Care and Use Committee (IACUC) at the SBP Medical Discovery Institute. Murine islets were isolated by collagenase P (Roche) perfusion as described following by histopaque-1077 (Sigma-Aldrich, Inc. St. Louis) gradient purification. Islets were handpicked and studied directly or after overnight culture in RPMI 1640 medium (Corning 10-040-CV) supplemented with 10% FBS, 1% penicillin/streptomycin, 100µg/ml primocin, 10mM Hepes, and 1mM sodium pyruvate.

### Validation of proinsulin interactions

For proinsulin interactions validated in human islets 2500 IEQ of normal human islets were lysed in 50mM Tris pH7.4, 150mM NaCl and 1% TX-100, 1X PMSF, 1X protease inhibitor cocktail and 1X phosphatase inhibitor cocktail (Thermo Scientific Pierce) on ice for 30 min. Lysates were precleared with protein A/G Agarose beads at 4°C for 1 hr before 132 µg of total protein extracts from human islet lysate with 132 µg of either proinsulin specific primary antibody (20G11) or mouse IgG Fc portion overnight at 4°C. Antibody bound proinsulin and its interactors were captured by binding to 100 µL of protein A agarose beads suspension 4 hrs (RT). Unbound supernatant was saved, and beads were washed with lysis buffer without detergent (1 × 100 µL and 1 × 500 µL) prior to eluting in 50 µL of 2X Laemmli buffer at 100°C for 10 min. 5% of either Lysate and Supernatant were run along with 12% of Elution (IP).

For proinsulin interactions validated in HEK293A cells, cells were plated in 6 or 12 well plates the day prior to transfection. Plasmids expressing human PRDX4-FLAG and Human QSOX1-HA were purchased from Sino Biological Inc. (Wayne, PA). Vectors expressing WT or Akita mutant c-Myc tagged hProins were obtained from Peter Arvan at University of Michigan (53). 2µg total DNA was transfected using Viafect transfection reagent following manufacturer instructions. PCDNA3.1 vector was used as the control and to match total amount DNA for co-transfection experiments. After 48 hours of transfection, cells were harvested, and protein was extracted with a lysis buffer on ice for 15-20 min. Supernatant was collected after spinning samples at 14000 rpm for 10 min at 4°C. Protein concentration was estimated by DC protein assay method (Biorad). Immunoprecipitation of tagged proinsulin, PRDX4 or QSOX1 was performed using agarose beads conjugated with anti-MYC, FLAG or HA respectively according to manufacturer protocol (Pierce). Freshly prepared lysates (250-350 µg total protein) were used for immunoprecipitation of c-Myc-Tag human proinsulin with c-Myc-Tag antibody coated magnetic beads using Magnetic c-Myc-Tag IP/Co-IP Kit (Pierce# 88844). Lysates were mixed with 25-50µl magnetic beads and incubated for 45 min at RT with mixing. After the incubation, supernatant was saved for analysis and after washing the beads, samples were eluted in 100µl of 1X non-reducing western blotting sample buffer. IP samples were reduced with DTT/ BME before separating onto the gel.

### Transfection of PRDX4-FLAG and Proinsulin-MYC to HEK293T cells

Transfection procedure was conducted similarly as described above for validation experiment except the following: (i) Lipofectamine transfection reagent (Thermo Scientific Pierce) was used according to manufacture recommended protocol, (ii) a combined amount of 2.5 µg of total DNA was utilized and, (iii) cells were lysed in RIPA buffer for subsequent immunoprecipitation, and 2X Laemmli without BME for subsequent SDS-PAGE analysis.

### Western blotting

Samples were prepared in Laemmli sample buffer without (non-reducing) or with (reducing) 2.5% BME. After boiling for 10 min at 100°C, samples were analyzed by SDS-PAGE (either 4-20% Tris-glycine or 4-12% Bis-tris pre-cast gel, Bio-rad Laboratories, Inc.) and transferred to nitrocellulose membranes (Bio-rad Laboratories, Inc.). Membranes were blocked with 5% BSA (4°C, 1hr) and cut according to molecular weight markers and incubated with corresponding primary antibodies (4°C, O.N): Rabbit BiP (for human islets), Rabbit GRP94 (Cell Signaling Tech, Danvers, MA), or Rabbit ERDJ5, Rabbit ERGIC1, Rabbit Myo18A and ERDJ3 (ProteinTech, Rosemont, IL), Goat PRDX4 (R&D System, Minneapolis, MN), Rabbit PRDX4 directly conjugated to Horse Radish Peroxidase (PRDX4-HRP) (LSBio, Seattle, WA), Mouse 20G11 (generated in house), Rabbit PRDX-SO_3_, Mouse HA and C-MYC (Abcam, Cambridge, MA). On murine islets, Rabbit vinculin (Proteintech, 66305-1-lg), Rabbit PDIA1 (Proteintech, 11245-1-AP), Mouse proinsulin (HyTest Ltd., 2PR8, CCI-17), and Rabbit BiP (for MIN6 cells, a kind gift from Dr Hendershot). For secondary antibodies, goat anti-mouse, goat anti-rabbit, donkey anti goat and donkey anti-guinea pig antibodies were used in 1:5000 (Li-Cor, IRDye®-800CW or IRDye®-680RD). After wash, membranes were imaged on Licor Odyssey CLX set up and western blot images were analyzed by ImageJ for quantitation of band intensity.

### PRDX4 knockdown

Viral vectors expressing lentiviral particles with 4 unique 29mer target-specific shRNA (A/B/C/D) to murine PRDX4 and 1 scramble control (non-specific or NS) were purchased from Origene (Rockville, MD). MIN6 cells were seeded on 48 well plates and cultured overnight in 0.3 mL medium. Polybrene (1mg/mL) was added at 1:125 ratio to culture medium before 20 µL of lentiviral particles (10^7^ TU/mL) was added to infect cells overnight. Infected cells were selected using Puromycin and subsequently by fluorescent activated cell sorting. For cell sorting, MIN6 cells were detached from plates using trypsin, quenched with FBS and resuspended in FACS buffer (1% FBS in PBS). Approximately 2 – 4 million cells were sorted for high GFP intensity, Cells were later transferred and maintained in MIN6 specific media (DMEM with 4.5g/L glucose and L-glutamine 3.4% NaHCO_3_, 1X penicillin, streptomycin, 275 nM BME, 15% FBS).

### qPCR analysis

Total RNAs were extracted from Wildtype MIN6 and MIN6 stably expressing shRNAs (non-specific/NS, C, D) using RNeasy kit (QIAGEN, Germantown, MD) and cDNAs were prepared using qScript cDNA SuperMix (Quantabio, Beverly, MA) according to manufacture recommended protocols. Primers used were: 18S (CCA GAG CGA AAG CAT TTG CCAAGA/ TCG GCA TCG TTT ATG GTC GGA ACT), CHOP (CTG CCT TTC ACC TTG GAG AC/ CGT TTC CTG GGG ATG AGA TA), PRDX4 (ACC AAG TAT TTC CCA CGA TAG TC/ GAT CAC TCC CTG CAT CTA AGC). The following method was used in a LightCycler 96 (Roche, Indianapolis, IN): 3 steps amplification (45 cycles) of 95°C/10s, 60°C/10s, 72°C, 10s followed by melting curve (1 cycle) 95°C/5s, 65°C/60s and cooling (1 cycle) 40°C/30s. Each gene was normalized by its corresponding 18S before quantification.

### Glucose stimulated insulin secretion (GSIS) protocol

Human islets were spun down briefly prior to washing with 1 × 20 mL and 1 × 5 mL PIMS complete media (PRODO labs, Aliso Viejo, CA). Islets were divided to aliquots and incubated in PIMS complete for approximately 4 hours at 37°C before starving with Glucose free media (Glucose free DMEM, 0.34% Sodium bicarbonate, sodium pyruvate, 15% FBS and penicillin/streptomycin) for 1 hr at 37°C. Glucose free media was removed and replaced with new Glucose free media supplemented with different glucose concentrations. After 18 hour incubation, media was removed and islets were washed with 1 × 1 mL PBS (with 20 mM NEM) before lysis with 2X Laemmli with 2 mM NEM and without BME. Similar GSIS protocol was used for MIN6.

### ELISA

Mouse insulin and ultra-sensitive human insulin ELISA kits were purchased from Mercodia (Uppsala, Sweden) and utilized according to manufacture recommended protocol.

## Acknowledgements

This work was supported by NIH R24 DK110973 (P I-A, PA and RJK), and JDRF research grant 2-SRA-2015-47-M-R (P I-A, RJK, WEB).

## Figure Legends

**Figure S1.**
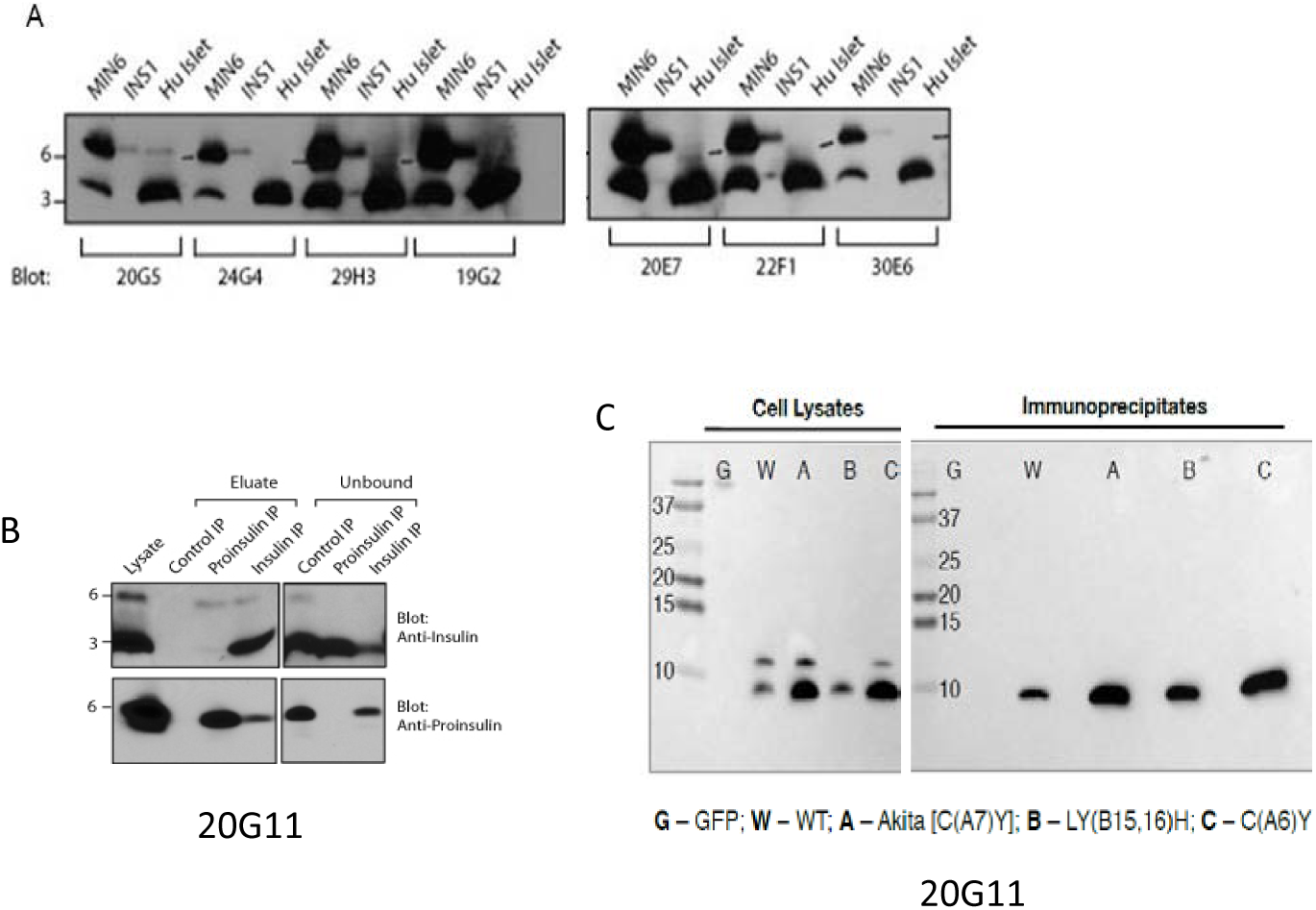
Antibody (20G11) Generation and Validation. Specificity of monoclonal antibodies to proinsulin. (A, B) Immunoblot of lysates from murine MIN6 cells, rat INS-1 cells and human islets with in-house generated monoclonal antibodies. C) WT proinsulin and proinsulin MIDY mutants were overexpressed in COS1 cells and immunoprecipitated with 20G11 antibody.

**Figure S2.**
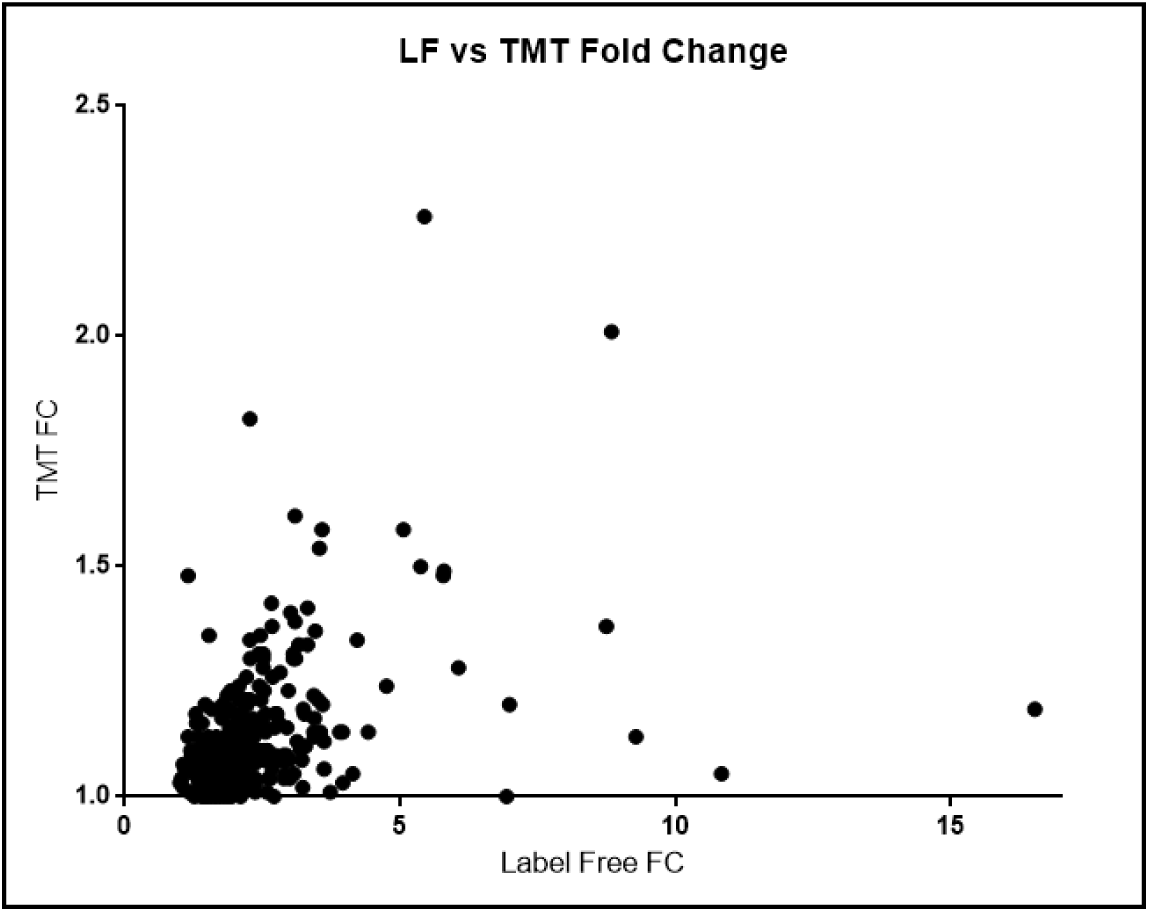
Comparison of Label Free and TMT Mass Spectrometric analysis on one islet preparation (HP-15157-01) identified common proteins but with different sensitivities.

**Figure S3.**
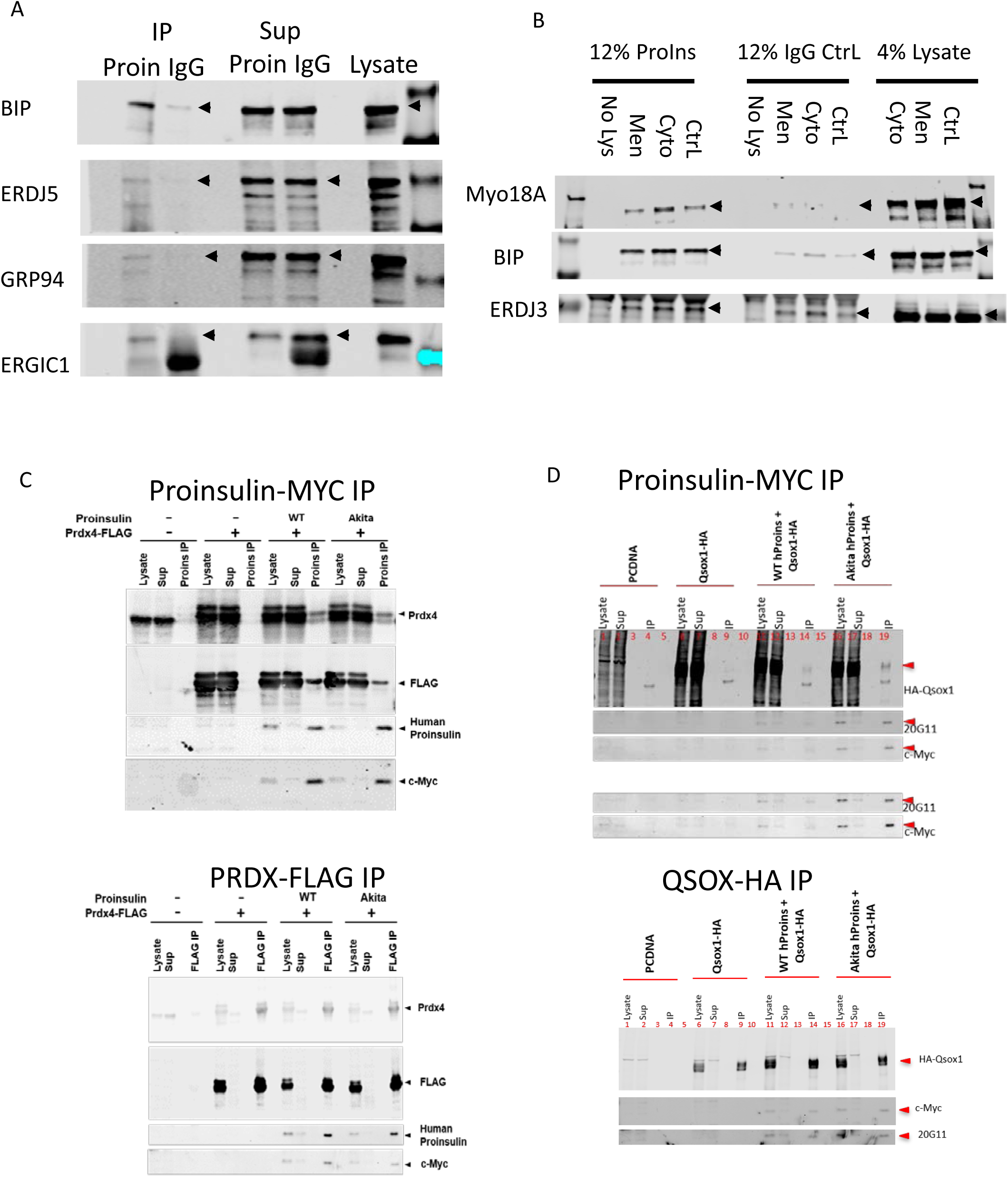
MS/MS Proinsulin Interactions Validated by AP-western blot. (A) Validation of BiP, ERDJ5, GRP94 and ERGIC1 interaction with proinsulin was performed by immunoprecipitation of from human islet lysates (HP-19038-01) with proinsulin specific primary antibody (20G11) or mouse IgG. 5% of Lysate and Supernatant (Sup) were run along with 12% of Elution (IP). Membranes were cut according to molecular weight markers and blotted for the indicated primary antibodies: Rabbit BIP, ERDJ5, GRP94 and ERGIC1. (B) Validation of Myo18A, BiP, and ERDJ3 interaction with proinsulin was performed by immunoprecipitation of lysates from a different human islet sample (HP-19038-01). 4% of lysate and 12% (IP) were analyzed on reducing gels. Membranes were cut according to molecular weight markers and blotted for the indicated primary antibodies: Rabbit Myo18A, BIP, ERDJ3. (C) Plasmids carrying human PRDX4-FLAG and/or hProins-c-Myc, and/ or pCDNA control plasmid were transiently transfected into HEK293 cells followed by immunoprecipitation with MYC-beads or FLAG beads. Lysates 5%, IPs 15% and supernatants 5% were blotted for PRDX4, FLAG, human proinsulin (20G11) and c-MYC. (D) Proinsulin Interaction with QSOX-1 and Myo18A. Plasmids carrying human QSOX1-HA and/or hProins-c-Myc or pCDNA control plasmid were transiently transfected into HEK293 cells followed by immunoprecipitation with MYC-beads (upper panel) or HA beads (lower panel). Lysates 5%, IPs 15% and supernatants 5% were blotted for HA (QSOX1) human proinsulin (20G11) and MYC tag (upper panel)

**Figure S4.**
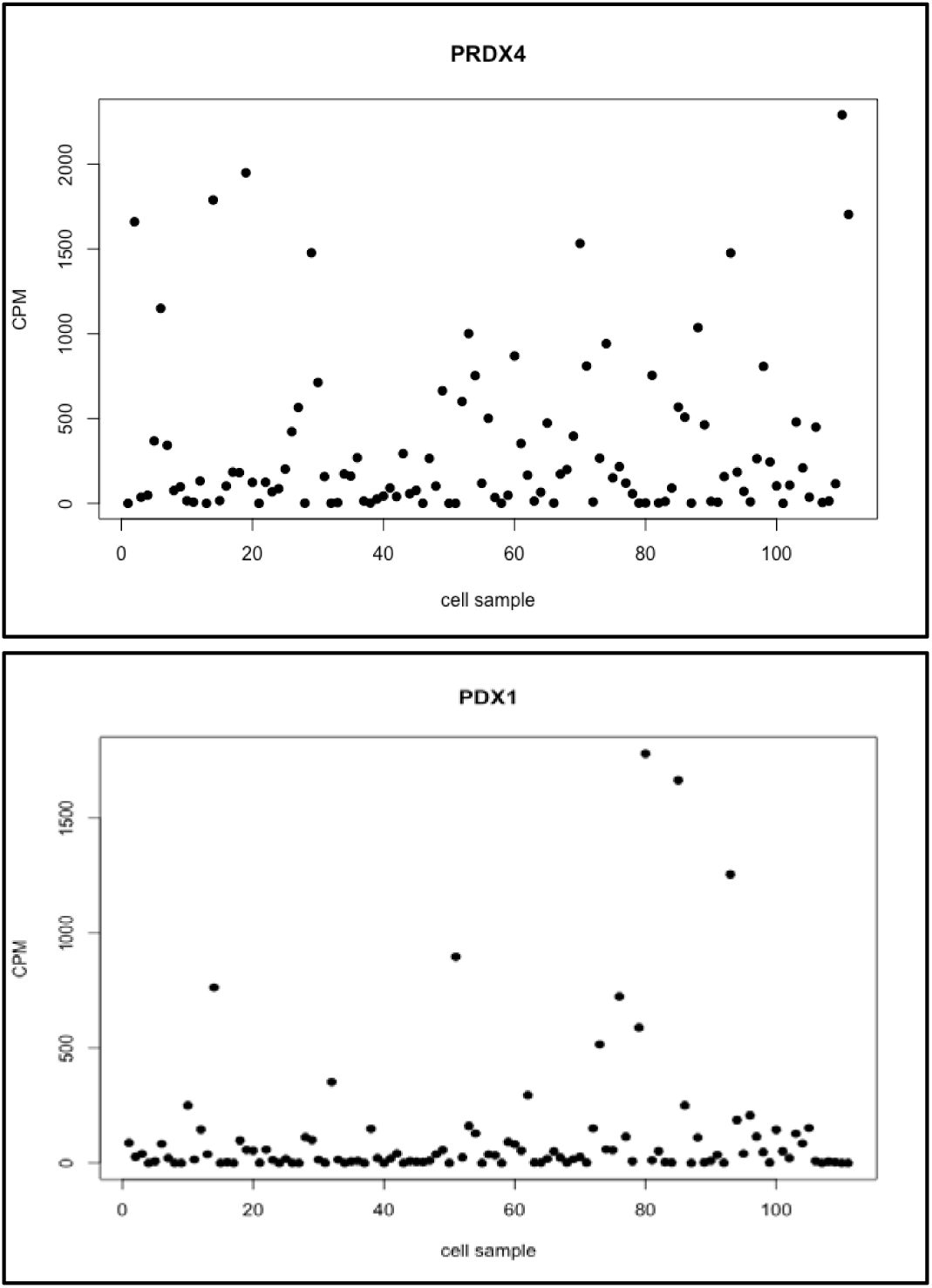
PRDX4 expression vs PDX1 expression in human beta cell single cell RNAseq (24).

**Figure S5.**
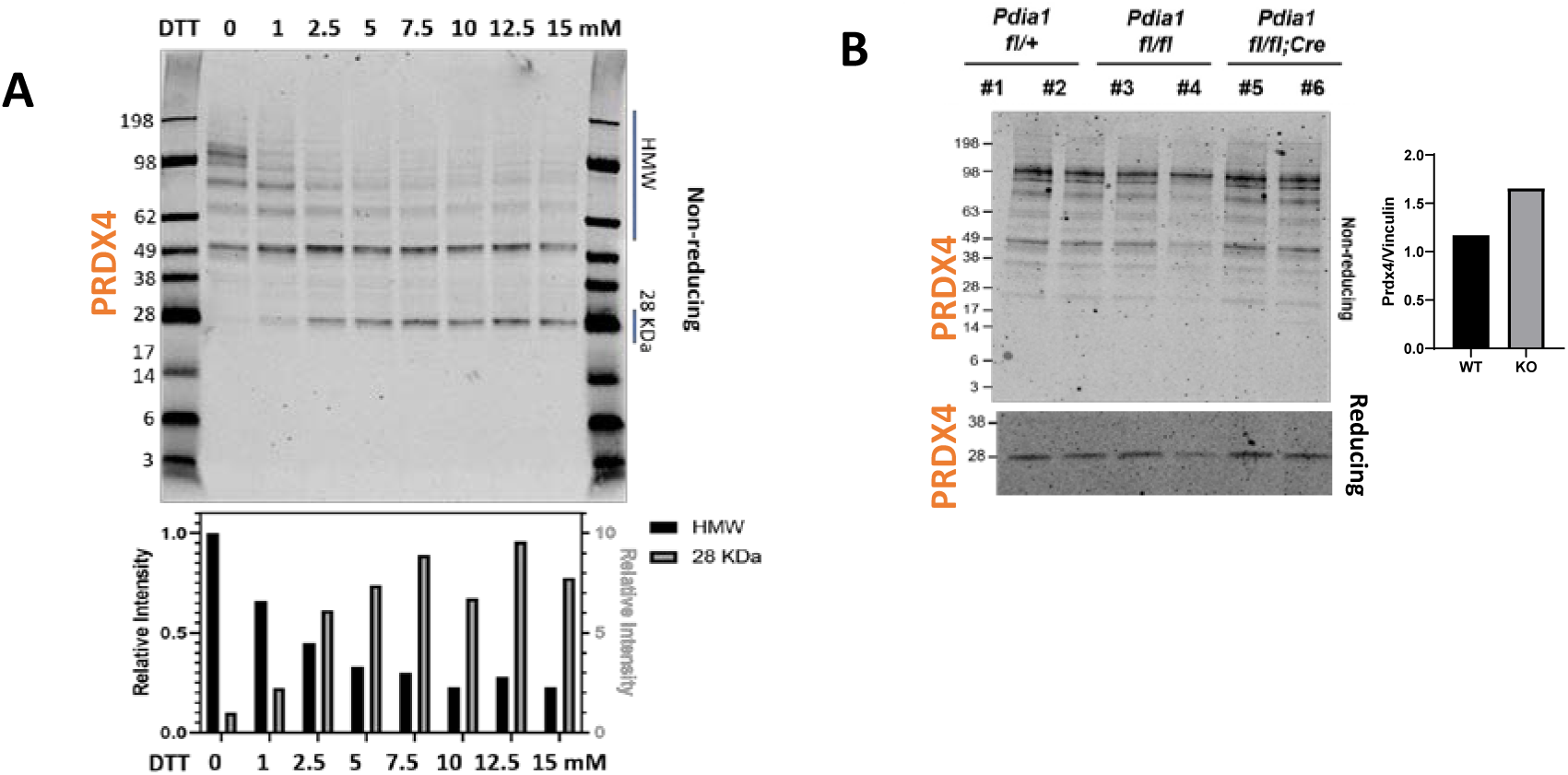
PRDX4 expression in islets from mice with beta cell specific PDIA1 deletion. (A) Lysates from untreated murine islets, untreated (0) or treated with a gradient of DTT concentrations ranging from 1 to 15 mM were run on a non-reducing gel and blotted for PRDX4. Quantification of disulfide linked HMW complex (size indicated 49 – 198 kDa) and PRDX4 monomer (28 kDa) intensity is graphed. (B) Islets from normal (PDIA1 fl/+ and PDIA1 fl/fl) mice versus PDI knockout female mice (PDIA1 fl/fl; Cre) were lysed, analyzed on non-reducing and reducing gels and blotted for PRDX4. Quantification of PRDX4 was performed by normalization to previously published vinculin (Reference (4) Fig 1C&D) as shown in accompanying graph (right).

**Figure S6.**
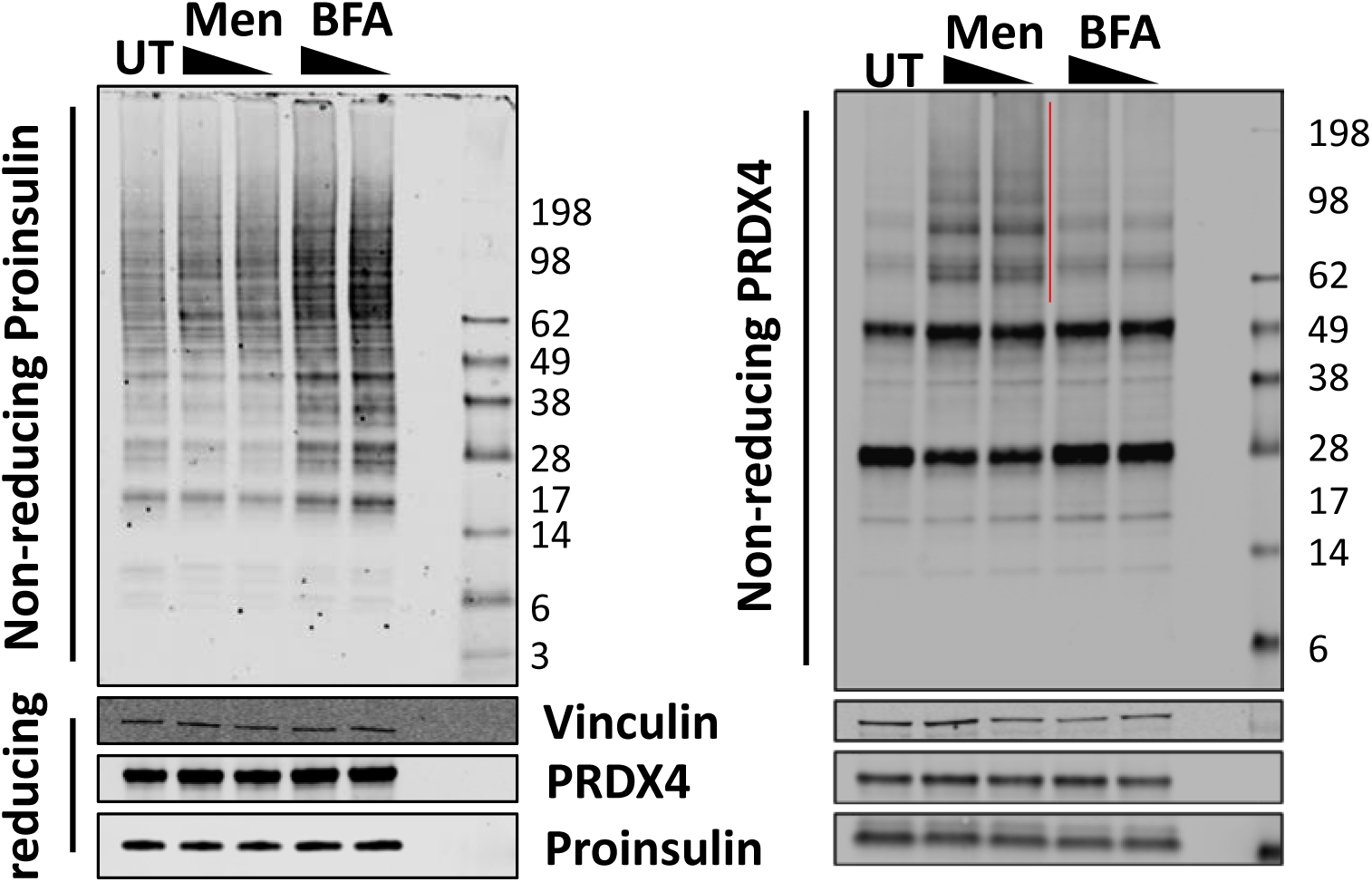
Menadione and BFA induce proinsulin misfolding and menadione recruits PRDX4 into HMW complexes in human islets. Human islets were treated with Menadione (200 and 100 µM, 1hr) and BFA (10 and 5 µg/mL, 1 hr).

## REFERENCES

1. Liu M, Weiss MA, Arunagiri A, Yong J, Rege N, Sun J, Haataja L, Kaufman RJ, Arvan P: Biosynthesis, structure, and folding of the insulin precursor protein. Diabetes, Obesity and Metabolism 2018;20:28–50

2. Schuit FC, Kiekens R, Pipeleers DG: Measuring the balance between insulin synthesis and insulin release. Biochemical and biophysical research communications 1991;178:1182–1187

3. Arunagiri A, Haataja L, Pottekat A, Pamenan F, Kim S, Zeltser LM, Paton AW, Paton JC, Tsai B, Itkin-Ansari P: Proinsulin misfolding is an early event in the progression to type 2 diabetes. eLife 2019;8:e44532

4. Jang I, Pottekat A, Poothong J, Yong J, Lagunas-Acosta J, Charbono A, Chen Z, Scheuner DL, Liu M, Itkin-Ansari P: PDIA1/P4HB is required for efficient proinsulin maturation and ß cell health in response to diet induced obesity. eLife 2019;8:e44528

5. Scheuner D, Kaufman RJ: The unfolded protein response: a pathway that links insulin demand with β-cell failure and diabetes. Endocrine reviews 2008;29:317–333

6. Lu M, Lawrence DA, Marsters S, Acosta-Alvear D, Kimmig P, Mendez AS, Paton AW, Paton JC, Walter P, Ashkenazi A: Opposing unfolded-protein-response signals converge on death receptor 5 to control apoptosis. Science 2014;345:98–101

7. Roth DM, Hutt DM, Tong J, Bouchecareih M, Page L, Wang N, Seeley T, Dekkers JF, Beekman JM, Garza D, Miller J, Masliah E, Morimoto RI, Balch WE: Modulation of the maladaptive stress response to mange diseases of protein folding. PloS Biol 2014;Submitted under final review

8. Back SH, Kaufman RJ: Endoplasmic reticulum stress and type 2 diabetes. Annual review of biochemistry 2012;81:767–793

9. Waanders LF, Chwalek K, Monetti M, Kumar C, Lammert E, Mann M: Quantitative proteomic analysis of single pancreatic islets. Proceedings of the National Academy of Sciences 2009;106:18902–18907

10. Ahmed M, Forsberg J, Bergsten P: Protein profiling of human pancreatic islets by two-dimensional gel electrophoresis and mass spectrometry. Journal of proteome research 2005;4:931–940

11. Schrimpe-Rutledge AC, Fontes G, Gritsenko MA, Norbeck AD, Anderson DJ, Waters KM, Adkins JN, Smith RD, Poitout V, Metz TO: Discovery of novel glucose-regulated proteins in isolated human pancreatic islets using LC-MS/MS-based proteomics. Journal of proteome research 2012;11:3520–3532

12. Zhang L, Lanzoni G, Battarra M, Inverardi L, Zhang Q: Proteomic profiling of human islets collected from frozen pancreata using laser capture microdissection. Journal of proteomics 2017;150:149–159

13. Liu M, Hodish I, Rhodes CJ, Arvan P: Proinsulin maturation, misfolding, and proteotoxicity. Proceedings of the National Academy of Sciences 2007;104:15841–15846

14. Megger DA, Pott LL, Ahrens M, Padden J, Bracht T, Kuhlmann K, Eisenacher M, Meyer HE, Sitek B: Comparison of label-free and label-based strategies for proteome analysis of hepatoma cell lines. Biochimica et Biophysica Acta (BBA)-Proteins and Proteomics 2014;1844:967–976

15. Choi M, Chang C-Y, Clough T, Broudy D, Killeen T, MacLean B, Vitek O: MSstats: an R package for statistical analysis of quantitative mass spectrometry-based proteomic experiments. Bioinformatics 2014;30:2524–2526

16. Wang YJ, Schug J, Won K-J, Liu C, Naji A, Avrahami D, Golson ML, Kaestner KH: Single-cell transcriptomics of the human endocrine pancreas. Diabetes 2016;65:3028–3038

17. Zito E, Melo EP, Yang Y, Wahlander Å, Neubert TA, Ron D: Oxidative protein folding by an endoplasmic reticulum-localized peroxiredoxin. Molecular cell 2010;40:787–797

18. Taft MH, Behrmann E, Munske-Weidemann L-C, Thiel C, Raunser S, Manstein DJ: Functional characterization of human myosin-18A and its interaction with F-actin and GOLPH3. Journal of Biological Chemistry 2013;288:30029–30041

19. Orci L, Ravazzola M, Perrelet A: (Pro)insulin associates with Golgi membranes of pancreatic B cells. PNAS 1984;81:6743–6746

20. Wu J, Kaufman R: From acute ER stress to physiological roles of the unfolded protein response. Cell death and differentiation 2006;13:374

21. Scheuner D, Vander Mierde D, Song B, Flamez D, Creemers JW, Tsukamoto K, Ribick M, Schuit FC, Kaufman RJ: Control of mRNA translation preserves endoplasmic reticulum function in beta cells and maintains glucose homeostasis. Nature medicine 2005;11:757

22. Tavender TJ, Springate JJ, Bulleid NJ: Recycling of peroxiredoxin IV provides a novel pathway for disulphide formation in the endoplasmic reticulum. The EMBO journal 2010;29:4185–4197

23. Rudolf J, Pringle MA, Bulleid NJ: Proteolytic processing of QSOX1A ensures efficient secretion of a potent disulfide catalyst. Biochemical Journal 2013;454:181–190

24. Wang YJ, Schug J, Won KJ, Liu C, Naji A, Avrahami D, Golson ML, Kaestner KH: Single-Cell Transcriptomics of the Human Endocrine Pancreas. Diabetes 2016;65:3028–3038

25. Paton AW, Beddoe T, Thorpe CM, Whisstock JC, Wilce MC, Rossjohn J, Talbot UM, Paton JC: AB 5 subtilase cytotoxin inactivates the endoplasmic reticulum chaperone BiP. Nature 2006;443:548

26. Pelham HR: Multiple targets for brefeldin A. Cell 1991;67:449–451

27. Klausner RD, Donaldson JG, Lippincott-Schwartz J: Brefeldin A: insights into the control of membrane traffic and organelle structure. The Journal of cell biology 1992;116:1071–1080

28. Hassler JR, Scheuner DL, Wang S, Han J, Kodali VK, Li P, Nguyen J, George JS, Davis C, Wu SP: The IRE1α/XBP1s pathway is essential for the glucose response and protection of β cells. PLoS biology 2015;13:e1002277

29. Shiu R, Pouyssegur J, Pastan I: Glucose depletion accounts for the induction of two transformation-sensitive membrane proteinsin Rous sarcoma virus-transformed chick embryo fibroblasts. Proceedings of the National Academy of Sciences 1977;74:3840–3844

30. Wood ZA, Schröder E, Robin Harris J, Poole LB: Structure, mechanism and regulation of peroxiredoxins. Trends in Biochemical Sciences 2003;28:32–40

31. Ghiasi SM, Dahlby T, Andersen CH, Haataja L, Petersen S, Omar-Hmeadi M, Yang M, Pihl C, Bresson SE, Khilji MS: Endoplasmic Reticulum Chaperone Glucose-Regulated Protein 94 Is Essential for Proinsulin Handling. Diabetes 2019;68:747–760

32. Pottekat A, Becker S, Spencer KR, Yates III JR, Manning G, Itkin-Ansari P, Balch WE: Insulin biosynthetic interaction network component, TMEM24, facilitates insulin reserve pool release. Cell reports 2013;4:921–930

33. Iuchi Y, Okada F, Tsunoda S, Kibe N, Shirasawa N, Ikawa M, Okabe M, Ikeda Y, Fujii J: Peroxiredoxin 4 knockout results in elevated spermatogenic cell death via oxidative stress. Biochemical Journal 2009;419:149–158

34. Caillard A, Sadoune M, Cescau A, Meddour M, Gandon M, Polidano E, Delcayre C, Da Silva K, Manivet P, Gomez A-M: QSOX1, a novel actor of cardiac protection upon acute stress in mice. Journal of molecular and cellular cardiology 2018;119:75–86

35. Day AM, Brown JD, Taylor SR, Rand JD, Morgan BA, Veal EA: Inactivation of a peroxiredoxin by hydrogen peroxide is critical for thioredoxin-mediated repair of oxidized proteins and cell survival. Molecular cell 2012;45:398–408

36. Ding Y, Yamada S, Wang K-Y, Shimajiri S, Guo X, Tanimoto A, Murata Y, Kitajima S, Watanabe T, Izumi H: Overexpression of peroxiredoxin 4 protects against high-dose streptozotocin-induced diabetes by suppressing oxidative stress and cytokines in transgenic mice. Antioxidants & redox signaling 2010;13:1477–1490

37. Lupi R, Del Guerra S, Mancarella R, Novelli M, Valgimigli L, Pedulli G, Paolini M, Soleti A, Filipponi F, Mosca F: Insulin secretion defects of human type 2 diabetic islets are corrected in vitro by a new reactive oxygen species scavenger. Diabetes & metabolism 2007;33:340–345

38. Ceriello A, Mercuri F, Quagliaro L, Assaloni R, Motz E, Tonutti L, Taboga C: Detection of nitrotyrosine in the diabetic plasma: evidence of oxidative stress. Diabetologia 2001;44:834–838

39. El Eter E, Al-Masri A: Peroxiredoxin isoforms are associated with cardiovascular risk factors in type 2 diabetes mellitus. Brazilian Journal of Medical and Biological Research 2015;48:465–469

40. Nabeshima A, Yamada S, Guo X, Tanimoto A, Wang KY, Shimajiri S, Kimura S, Tasaki T, Noguchi H, Kitada S, Watanabe T, Fujii J, Kohno K, Sasaguri Y: Peroxiredoxin 4 protects against nonalcoholic steatohepatitis and type 2 diabetes in a nongenetic mouse model. Antioxid Redox Signal 2013;19:1983–1998

41. Hutt DM, Balch WE: Expanding proteostasis by membrane trafficking networks. Cold Spring Harbor perspectives in biology 2013;5:a013383

42. Powers ET, Balch WE: Diversity in the origins of proteostasis networks—a driver for protein function in evolution. Nature reviews Molecular cell biology 2013;14:237

43. Chawade A, Alexandersson E, Levander F: Normalyzer: a tool for rapid evaluation of normalization methods for omics data sets. Journal of proteome research 2014;13:3114–3120

44. Clough T, Thaminy S, Ragg S, Aebersold R, Vitek O: Statistical protein quantification and significance analysis in label-free LC-MS experiments with complex designs. BMC bioinformatics 2012;13:S6

45. Tripathi S, Pohl MO, Zhou Y, Rodriguez-Frandsen A, Wang G, Stein DA, Moulton HM, DeJesus P, Che J, Mulder LC: Meta-and orthogonal integration of influenza “OMICs” data defines a role for UBR4 in virus budding. Cell host & microbe 2015;18:723–735

46. Kuleshov MV, Jones MR, Rouillard AD, Fernandez NF, Duan Q, Wang Z, Koplev S, Jenkins SL, Jagodnik KM, Lachmann A: Enrichr: a comprehensive gene set enrichment analysis web server 2016 update. Nucleic acids research 2016;44:W90–W97

47. Uhlén M, Fagerberg L, Hallström BM, Lindskog C, Oksvold P, Mardinoglu A, Sivertsson Å, Kampf C, Sjöstedt E, Asplund A: Tissue-based map of the human proteome. Science 2015;347:1260419

48. Alanis-Lobato G, Andrade-Navarro MA, Schaefer MH: HIPPIE v2. 0: enhancing meaningfulness and reliability of protein–protein interaction networks. Nucleic acids research 2016:gkw985

49. Vanunu O, Magger O, Ruppin E, Shlomi T, Sharan R: Associating genes and protein complexes with disease via network propagation. PLoS computational biology 2010;6:e1000641

50. Huang S-sC, Fraenkel E: Integrating proteomic, transcriptional, and interactome data reveals hidden components of signaling and regulatory networks. Sci Signal 2009;2:ra40–ra40

51. Loguercio S: Network-Augmented Genomic Analysis (NAGA) applied to cystic fibrosis studies. F1000Research 2015;4

52. Shannon P, Markiel A, Ozier O, Baliga NS, Wang JT, Ramage D, Amin N, Schwikowski B, Ideker T: Cytoscape: a software environment for integrated models of biomolecular interaction networks. Genome research 2003;13:2498–2504

53. Liu M, Haataja L, Wright J, Wickramasinghe NP, Hua Q-X, Phillips NF, Barbetti F, Weiss MA, Arvan P: Mutant INS-gene induced diabetes of youth: proinsulin cysteine residues impose dominant-negative inhibition on wild-type proinsulin transport. PloS one 2010;5:e13333

